# Systems-level investigation of mucopolysaccharidosis IIIA identifies deficient synaptic activity as a key driver of disease progression

**DOI:** 10.1101/2022.10.03.510585

**Authors:** Alon M Douek, Abbas Salavaty, Florian Kreuder, Sebastian-Alexander Stamatis, Joel R Steele, Iresha Hanchapola, Anup D Shah, Ralf B Schittenhelm, Mirana Ramialison, Peter D Currie, Jan Kaslin

## Abstract

Mucopolysaccharidoses are lysosomal storage diseases that collectively represent a major cause of lethal, treatment-refractory childhood dementias ^1–7^ Clinically-useful interventions are hampered due to an incomplete understanding of their neuropathological origins. Using the zebrafish *sgsh* model of mucopolysaccharidosis IIIA ^8^ (MPS IIIA, Sanfilippo syndrome A), we conducted several ‘omics-based analyses, and developed and benchmarked a novel bioinformatic feature classification and ranking model for high-throughput datasets – ExIR – to prioritise important features in the progression of neurological manifestations of the disease. We find that the massive endolysosomal burden resulting from increased lysosomal storage of heparan sulfate and other secondarily accumulating substrates, such as sphingolipids, induces abnormal microtubule organisation and vesicle trafficking in neurons. This results in a gradual impairment of synaptic vesicle localisation at the presynaptic terminal and consequently impaired neuronal activity. Importantly, the endolysosomal phenotype in MPS IIIA zebrafish well-precedes the onset of neural pathology, though the larval MPS IIIA brain was found to be more susceptible to perturbation than wild type siblings. Collectively, these analyses demonstrate the presence of a progressive ‘functional neurodegenerative’ phenotype underpinning neurological disease in MPS IIIA. Our findings provide direct mechanistic evidence linking the well-described lysosomal storage basis for MPS IIIA to its disproportionately severe neural clinical involvement, enabling development and refinement of future therapeutic interventions for this currently untreatable disorder.

**Highlights:**

- MPS IIIA represents one of the most common causes of broadly fatal childhood dementia, but the mechanisms underlying disease progression are poorly understood.
- The first systems-level analyses of disease state and progression in the CNS of an MPS IIIA animal model were performed.
- **Ex**perimental data-based **I**ntegrative **R**anking (ExIR) was developed to provide unbiased prioritisation and classification of biological data as drivers, biomarkers and mediators of biological processes from high-throughput data at a systems level.
- Application of ExIR to a transcriptomic and proteomic analyses of a zebrafish model of MPS IIIA implies progressive deficiencies in synaptic activity as a key driver of disease progression correlating with progressive neuronal endolysosomal burden and secondary storage diseases.
- A novel unifying explanation of pathobiology and progression of MPS IIIA facilitates identification of clinically targetable features and may be generalised to other neuronopathic storage disorders.

## Introduction

Mucopolysaccharidosis IIIA (MPS IIIA, Sanfilippo syndrome A, OMIM #252900) is a paediatric neurodegenerative disease resulting from impaired catabolism of the glycosaminoglycan heparan sulfate (HS) due to loss of enzymatic function in the lysosomal sulfoglucosamine sulfohydrolase *SGSH.* Though this loss of function results in global accumulation of HS, MPS IIIA patients primarily present clinically with central nervous system (CNS)-specific symptoms, which represent the primary source of morbidity. It remains unclear why the CNS is particularly adversely affected in this disease state despite global HS accumulation, and the specific mechanisms and drivers of CNS disease progression are similarly incompletely understood. As the neuronopathic and cognitive features of MPS IIIA are similar to those observed in other mucopolysaccharidoses ^5^, a generalisable mechanism underlying these pathologies would be highly clinically relevant and may inform future work towards therapies. In order to better investigate and molecularly dissect MPS IIIA CNS pathology, we previously generated and characterised the first zebrafish model of MPS IIIA ^8^, utilising the excellent genetic tractability of the zebrafish ^9^ to demonstrate highly conserved pathological features between the model and human MPS IIIA. We sought to further explore the molecular basis of MPS IIIA CNS disease progression through the first transcriptomic and proteomic analyses of the whole CNS of an MPS IIIA disease model. To robustly determine transcriptomic features underlying disease progression in the CNS of MPS IIIA zebrafish in an unbiased manner, we developed ExIR (**<u>Ex</u>**perimental data-based **<u>I</u>**ntegrative **<u>R</u>**anking), a novel feature classification and prioritisation model that accurately and sensitively identified several drivers which collectively provide a novel mechanistic framework for the basis of CNS pathology in MPS IIIA. Unlike existing prioritisation models which rely on external data sources (e.g. gene/pathway ontology and protein-protein interaction databases) or the presence of a biological ‘ground truth’ to rank features, ExIR is a standalone model which extracts, classifies and prioritises candidate features *(e.g.* genes, proteins etc.) solely from high-throughput experimental data. Benchmarking using several different transcriptomic and proteomic datasets demonstrated that ExIR outperforms other prioritisation methods in specifically and sensitively identifying, classifying and ranking features by their functional importance. By utilising ExIR-prioritised candidate genes from brain-wide transcriptomic and proteomic profiling of the MPS IIIA zebrafish, we provide a wholistic model of MPS IIIA CNS pathology implicating cumulative, microtubule-dependent neuronal endolysosomal storage of HS as well as other secondary substrates. Subsequent impaired intracellular trafficking of neurotransmitter-containing vesicles depleted the presynaptic reserve pool, leading to progressively-worsening defective synaptic activity and driving functional neurodegeneration in MPS IIIA.

### Transcriptomic and proteomic profiling of early and late CNS-specific features of MPS IIIA zebrafish

A major outstanding issue limiting development of effective clinical interventions for MPS IIIA is a lack of understanding of the fundamental mechanisms underlying disease progression. While it is known that MPS IIIA results from accumulation of HS due to *SGSH* dysfunction, it remains unclear how this leads to the manifestation of a functional neurodegenerative phenotype. While several animal models of MPS IIIA exist, none so far have been utilised for CNS-specific systems-level analyses. To this end, we performed bulk RNA sequencing of young (3-month-old) and aged (18-month old) brains from the homozygous *sgsh^Δex5-6^* zebrafish mutant ^8^ (aka *sgsh^mnu301^,* hereafter referred to as *sgsh)* and age-matched wild type sibling controls towards better understanding the biological processes and mechanisms underlying MPS IIIA and its CNS pathology.

Between young wild type and *sgsh* brains, 26 genes were differentially expressed (DE, 12 up, 14 down, **Fig. 1a**). Most downregulated DEGs in young *sgsh* brains were immediate-early genes (IEGs)/primary response genes ^10^, namely *npas4a, fosab, egr1, egr2a, egr4, btg2, junba, junbb, nr4a1, ier2a* and *npas4l*. These IEGs represent a class of transcription factors that – in the context of the CNS – are induced in postsynaptic neurons following synaptic neurotransmission, where they are able to direct downstream neural circuit activity in a MAPK-dependent, *de novo* translation-independent manner ^10,11^. Upregulated DEGs in young *sgsh* brain included *tyrp1b, pmela* and *pmelb* (components of the premelanosome, a lysosome-like organelle ^12^), and *baiap2b,* involved in membrane trafficking and cytoskeletal organisation ^13^ as well as a component of CNS synapses ^14^. Examination of enriched gene ontology biological processes (GO-BPs) in DEGs in the young *sgsh* brain demonstrate overrepresentation of transcription factor-associated functions associated with downregulation of multiple IEGs (**Fig. 1b**). Analysis of potential protein-protein interactions between young brain DEGs using STRING ^15^ demonstrates two major clusters of DEGs; one comprising the downregulating IEGs, and the other associated with components of premelanosome assembly (**Fig. 1c**).

**Fig. 1.**
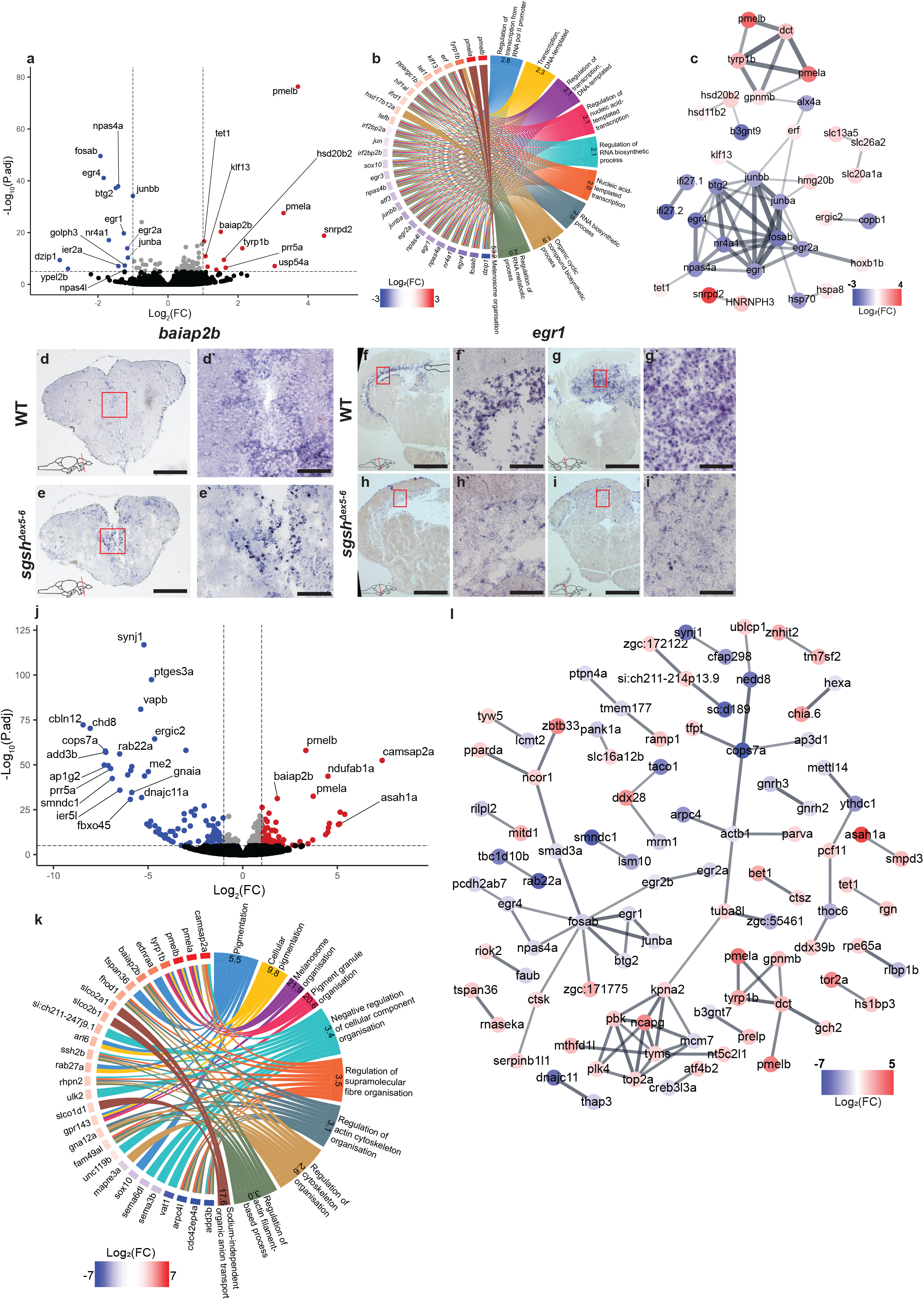
Transcriptomic analyses of young and aged *sgsh* zebrafish brain identifies profound downregulation of immediate early genes (IEGs). **a,** Volcano plot of differentially-expressed genes between young (3-month-old) wild type and *sgsh* brain by RNA sequencing. **b,** Chord plot of relationship between top 10 GO-BPs enriched in the young cohort, and their association to young brain DEGs; term fold enrichment given in each chord. **c,** Protein-protein interaction (PPI) network of young DEGs. Nodes coloured by log_2_(FC). **d-d’,** Expression of *baiap2b* in wild type telencephalon detected by *in situ* hybridisation (ISH). Scale bar 200 μm in **d** and 40 μm in **d’**. **e-e’** Expression of *baiap2b* in *sgsh* homozygous telencephalon detected by ISH. Scale bar 200 μm in **e** and 40 μm in **e’**. **f-f’**, *egr1* expression by ISH in wild type optic tectum. Scale bar 300 μm in **f** and 50 μm in **f’**. **g-g’,** *egr1* expression by ISH in wild type cerebellum. Scale bar 300 μm in **g** and 50 μm in **g’**. **h-h’**,*egr1* expression by ISH in *sgsh* homozygous optic tectum. Scale bar 300 μm in **h** and 50 μm in **h’**. **i-i’,** *egr1* expression by ISH in *sgsh* homozygous cerebellum. Scale bar 300 μm in **i** and 50 μm in **i’**. **j,** Volcano plot of DEGs between aged (18-month-old) wild type and *sgsh* brain by RNA sequencing. **k,** Chord plot of relationship between top 10 GO-BPs enriched in the aged cohort, and their association to aged brain DEGs; term fold enrichment given in each chord. **l,** PPI network of aged DEGs. Nodes coloured by log_2_(FC).

We next sought to determine if the differential gene expression patterns observed by RNA sequencing were associated with specific neuroanatomical domains. *In situ* hybridisation (ISH) for *baiap2b* in adult wild type and *sgsh* brain demonstrated roughly equivalent levels of expression in neuronal nuclei in the dorsal tegmentum and cells in the cerebellar Purkinje layer in both wild type and *sgsh* brains (**Extended Data Fig. 1a-d’**). While weak expression was observed in telencephalic neurons in wild type brains (**Fig. 1d-d’**), *baiap2b* was strongly expressed in this domain in homozygous *sgsh* siblings (**Fig. 1e-e’**). Contrastingly, *egr1* was observed to be robustly expressed in the optic tectum periventricular grey zone (**Fig. 1f-f’**) and cerebellar granule cell layer in wild type brain (**Fig. 1g-g’**), but almost completely abolished from these regions in *sgsh* brains (**Fig. 1h-h’, i-i’**).

A much greater number of DEGs were detected between aged *sgsh* and wild type brains compared to young samples (166 genes; 78 up, 88 down, **Fig. 1j**), indicative of a significantly perturbed brain transcriptome associated with the progression of CNS disease. Eleven genes were consistently DE across both young and aged timepoints (42.31% of young DEGs, 6.63% of aged DEGs – namely, *pmela, pmelb, tyrp1b, prr5a, baiap2b, tet1, junba, egr2a, egr1, npas4a* and *fosab*), and thus are likely to be most fundamentally involved in CNS pathology in the *sgsh* zebrafish – the direction of differential expression was the same over time for all genes except *prr5a*, which is upregulated in young *sgsh* brain compared to wild type siblings but downregulated in aged *sgsh* brains. Aside from these temporally conserved DEGs, a large proportion of downregulated genes in aged *sgsh* brains indicated involvement in membrane and endo-lysosomal trafficking *(synj1, rab22a, vapb, ergic2, ap1g2),* protein aggregate degradation *(ptges3a, dnajclla, cops7a, fbxo45),* and cytoskeletal remodelling *(add3b)*. In addition to *pmela, pmelb* and *baiap2b,* the non-centrosomal microtubule organiser *camsap2a*, the acid ceramidase *asah1a* and the mitochondrial respiratory complex subunit *ndufab1a* were upregulated in aged *sgsh* brains. GO-BP analysis demonstrated enrichment of melanosome-associated genes, in addition to hallmarks of endo-lysosomal trafficking and associated cytoskeletal remodelling (**Fig. 1k**). STRING network analysis highlighted two major node clusters; one comprising multiple IEGs and centring on *fosab*, and another primarily composed of up-regulated genes associated with mitosis or nucleotide metabolism (*kpna2*, *pbk*, *ncapg*, *tyms, top2a, plk4* – only one gene, *mcm7*, in this cluster is down-regulated) (**Fig. 1l**). Other smaller gene clusters relate to the pigmentation-associated genes detected as DE in young *sgsh* brains *(pmela, pmelb, tyrp1b, dct,* **Fig. 1l**). Taken together, transcriptomic analyses of young and aged *sgsh* brains highlight early neuronal activity-related deficits preceding subsequent, wide-scale cellular dysfunction in MPS IIIA.

While the above transcriptomic analyses are sensitive in detecting acute patterns of gene expression associated with the underlying neuropathological features of MPS IIIA, they are less robust in highlighting chronic features that accumulate in the CNS during disease progression. Such features are better explained by examination of the translational output of cells in affected tissues, whose latency within a cell does not necessarily correlate to the timing of gene expression. To define the translational signature of young and aged *sgsh* brains, we performed quantitative TMT (tandem mass tag) label-based mass spectrometry (nanoLC-ESI MS/MS) on age-matched wild type or *sgsh* brains at the same time points as the previous transcriptomic analyses. Thirty-three proteins were found to be differentially abundant (DA) between young wild type and *sgsh* brains; 30 of these proteins were enriched in *sgsh* brains, while only three were enriched in wild type brains (**Fig. 2a**). Protein-protein interaction (PPI) analysis with STRING showed strong interactivity and functional relationships between most DA proteins. The majority of DA proteins (DAPs) were either lysosomal proteases/peptidases including multiple cathepsins (Ctsh, Ctsla, Ctsba and Loc100333521 [designated here as Ctsz-l based on orthology]), or were associated with lysosomal degradation of glycosaminoglycans, gangliosides or other sphingolipids (Gnsa, Gusb, Hexb, Naga, Gaa2) (**Fig. 2b**). This was supported by pathway, process and organellar enrichment analyses demonstrating that most DAPs were directly associated with lysosomal catabolic functions (**Fig. 2c**). Similarly, 38 proteins were differentially abundant between aged brains from both genotypes, with 35 enriched in the aged *sgsh* brain, and three enriched in the brain of wild type siblings (**Fig. 2d**). As in young *sgsh* brains, most DAPs in the aged *sgsh* brain were directly associated with protease activity or lysosomal substrate degradation (**Fig. 2e-f**). Notably, only one protein – the arylsulfatase Arsa – was consistently deficient in both young and aged *sgsh* brains compared to wild type brains. Human ARSA deficiency causes metachromatic leukodystrophy ^16^ (OMIM #250100), a lysosomal sphingolipid storage disease with several overlapping clinical features with MPS IIIA. The proteomic data strongly suggested the presence of both lysosomal accumulation as well as secondary sphingolipid storage; indeed, LAMP1 immunoreactivity and LipidSpot 610 staining demonstrated significantly increased lipid storage in the dense interneuron populations of the midbrain (**Fig. 2g, h**) and cerebellum (**Fig. 2i, j**) in adult *sgsh* brain compared to wild type siblings. In the wild type midbrain, large cytoplasmic LAMP1^+^ organelles (interpreted here as endolysosomes) were observed predominantly in the ventricular radial glial population, many of which stained positive for lipid storage using the neutral lipid dye LipidSpot 610 (**Fig. 2g**). In contrast, massively increased LAMP1 immunoreactivity was seen in more dorsal neuronal layers in the *sgsh* tectum (**Fig. 2h**). LAMP1 immunoreactivity was sparse in the cerebellar granule neuron layer of wild type siblings (**Fig. 2i and boxed region**), but was strongly detected in putative microglia throughout the molecular layer of the *sgsh* cerebellum (**Fig. 2j and boxed region**) supporting our previous observations of profound neuroinflammation in the *sgsh* zebrafish brain ^8^. Additionally, large, electron-dense lipid-containing organelles were routinely observed (**Fig. 2m**) across the *sgsh* CNS, alongside extremely dense networks of axonal microtubules (**Fig. 2n, n’**) compared to those of wild type siblings (**Fig. 2l, l’**). These observations suggest an increased requirement for trafficking of heparan sulfate- and lipid-containing endolysosomes in the MPS IIIA zebrafish CNS. Of note, LAMP1 immunoreactivity was significantly reduced in wild type (7 dpf) larval brains (**Fig. 2o**) compared to homozygous *sgsh* siblings (**Fig. 2p**), indicating the increased endolysosomal burden in the *sgsh* mutant is an early feature of the protopathological state. This finding supports our previous observation of lysosomal accumulation in larval *sgsh* zebrafish brains using the LysoTracker dye ^8^. Taken together, proteomics analyses of young and aged brains highlight consistent, massive endolysosomal burden in the *sgsh* CNS due to increased intracellular storage of diverse content, including lipids and partially-degraded GAG catabolites.

**Fig. 2.**
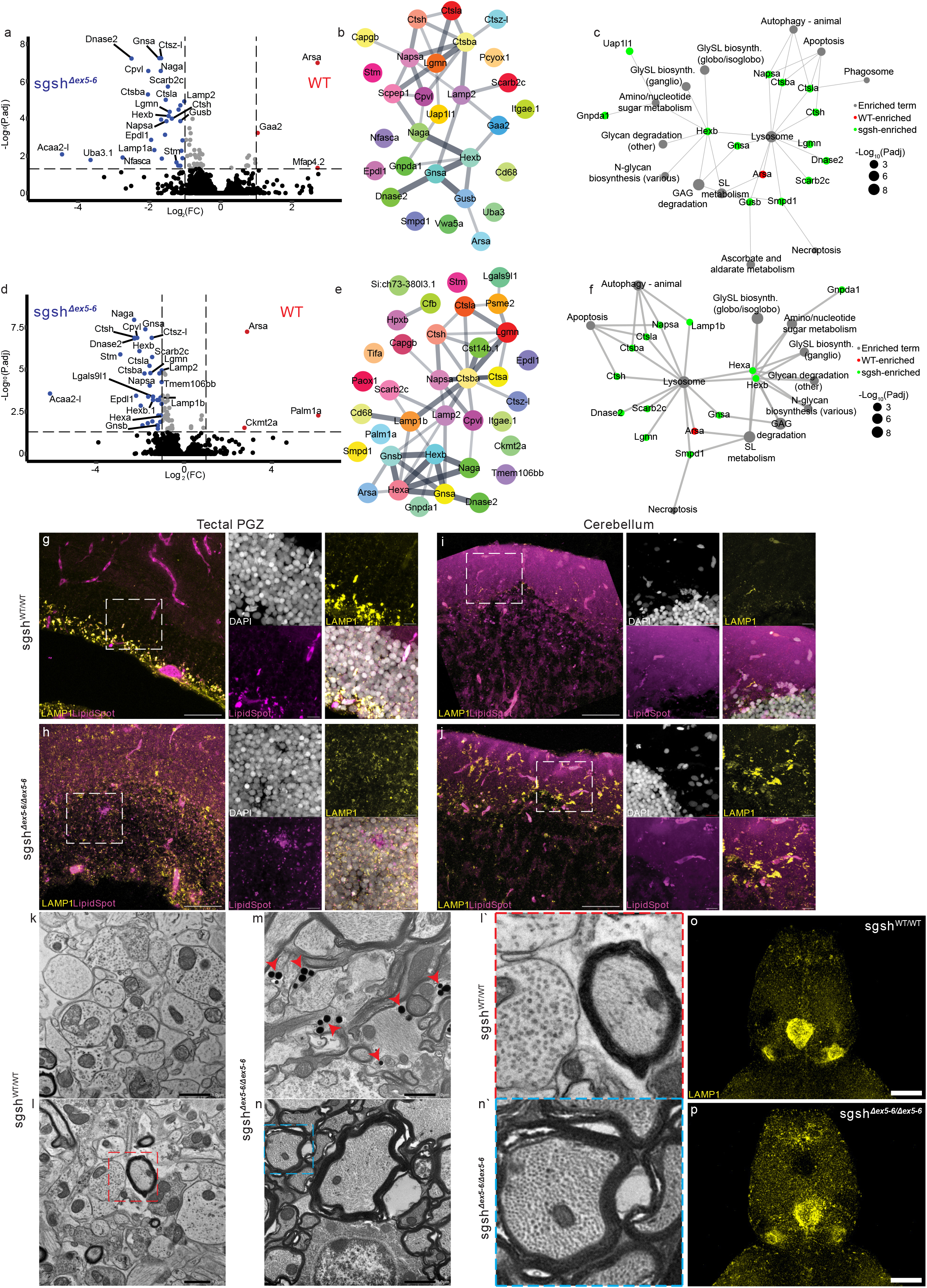
Quantitative protein analyses of young and aged *sgsh* zebrafish brain highlights compensatory lysosomal accumulation associated with primary and secondary substrate degradation. **a,** Volcano plot of differentially-abundant proteins (DAPs) between young (3-month-old) wild type and *sgsh* brain by quantitative (TMT label-based) mass spectrometry. **b,** STRING PPI network of young brain DAPs. Edge weight represents confidence of proteinprotein association. **c,** Term-feature graph of young brain DAPs and associated KEGG pathways. Red protein nodes enriched in WT brain; green protein nodes enriched in *sgsh* brain. Grey KEGG pathway node size reflects degree of statistical significance of enrichment. **d,** Volcano plot of differentially-abundant proteins (DAPs) between aged (18-month-old) wild type and *sgsh* brain by quantitative (TMT label-based) mass spectrometry. **e,** STRING PPI network of aged brain DAPs. Edge weight represents confidence of protein-protein association. **f,** Term-feature graph of aged brain DAPs and associated KEGG pathways. Red protein nodes enriched in WT brain; green protein nodes enriched in *sgsh* brain. Grey KEGG pathway node size reflects degree of statistical significance of enrichment. **g,** LAMP1 immunoreactivity and LipidSpot 610 lipid droplet staining in wild type optic tectum; scale bar 50 μm. Boxed region displayed in panels, scale bar 10 μm. **h,** LAMP1 immunoreactivity and LipidSpot 610 lipid droplet staining in *sgsh* optic tectum; scale bar 50 μm. Boxed region displayed in panels, scale bar 10 μm. **i,** LAMP1 immunoreactivity and LipidSpot 610 lipid droplet staining in wild type cerebellum; scale bar 50 μm. Boxed region displayed in panels, scale bar 10 μm. **j,** LAMP1 immunoreactivity and LipidSpot 610 lipid droplet staining in *sgsh* cerebellum; scale bar 50 μm. Boxed region displayed in panels, scale bar 10 μm. **k-l’,** TEM of wild type brain; scale bar 1 μm in **k-l**. **l’** is boxed region in **l**. **m-n’,** TEM of *sgsh* brain; scale bar 1 μm in **m-n**. **n’** is boxed region in **n**. Red arrowheads in **m** are enlarged lipid droplets. **o,** LAMP1 immunoreactivity in wild type 7 dpf brain. LAMP1 expression is generally weakly ubiquitous, with concentrated expression in the choroid plexus and habenula. Scale bar 50 μm. **p,** LAMP1 immunoreactivity in *sgsh* 7 dpf brain. Note strong, dispersed LAMP1 expression throughout the brain parenchyma. Scale bar 50 μm.

### Development and benchmarking of the ExIR workflow

Systems-level analyses of the *sgsh* zebrafish brain demonstrated the presence of complex transcriptional and protein signatures of MPS IIIA progression in our model. In particular, the drastic increase in the number of DEGs detected between young and aged wild type vs. *sgsh* brain transcriptomes highlights this complexity, wherein disease progression is reflected in significant derangement of homeostatic patterns of gene expression. In such perturbed states, it can be difficult to determine from these large gene/protein lists the most important features that are likely to be most relevant to the initiation and progression of the disease state, rather than secondary or indirect consequences of the pathobiology. Additionally, guided analyses of high-throughput data often rely on existing ontological information and manual annotations of gene/protein function. As such knowledge sources are derived from the broader literature, they are inherently in a state of flux; despite their effectiveness at communicating categorical information, these ontologies may fail to accurately account for heterogeneity or complexity inherent to the analysed data ^17^. In many cases, including the systems-level analyses of the *sgsh* zebrafish CNS, it would be desirable to determine importance of features in high-throughput datasets without relying on external sources of information.

To this end, we established the Experimental data-based Integrative Ranking (ExIR) model, which solely utilises high-throughput experimental data to calculate several scores used to classify and rank features from that data (**Fig. 3a-b, supplementary methods**). In ExIR, normalised high-throughput data is input, and an adjacency matrix is generated through correlation analysis. Correlated features are then used to generate a final matrix used in network reconstruction. On the basis of combinations of the derived scores (**Fig. 3b**), features are categorised as drivers, biomarkers and/or mediators prior to ranking features within each class. This is performed in two major steps, first employing supervised machine learning (ML) through a random forest algorithm to filter the input normalised data, then unsupervised ML by way of principal component analysis to generate a rotation value for network assembly (**Fig. 3a**, further detailed in **Supplementary Methods**).

**Fig. 3.**
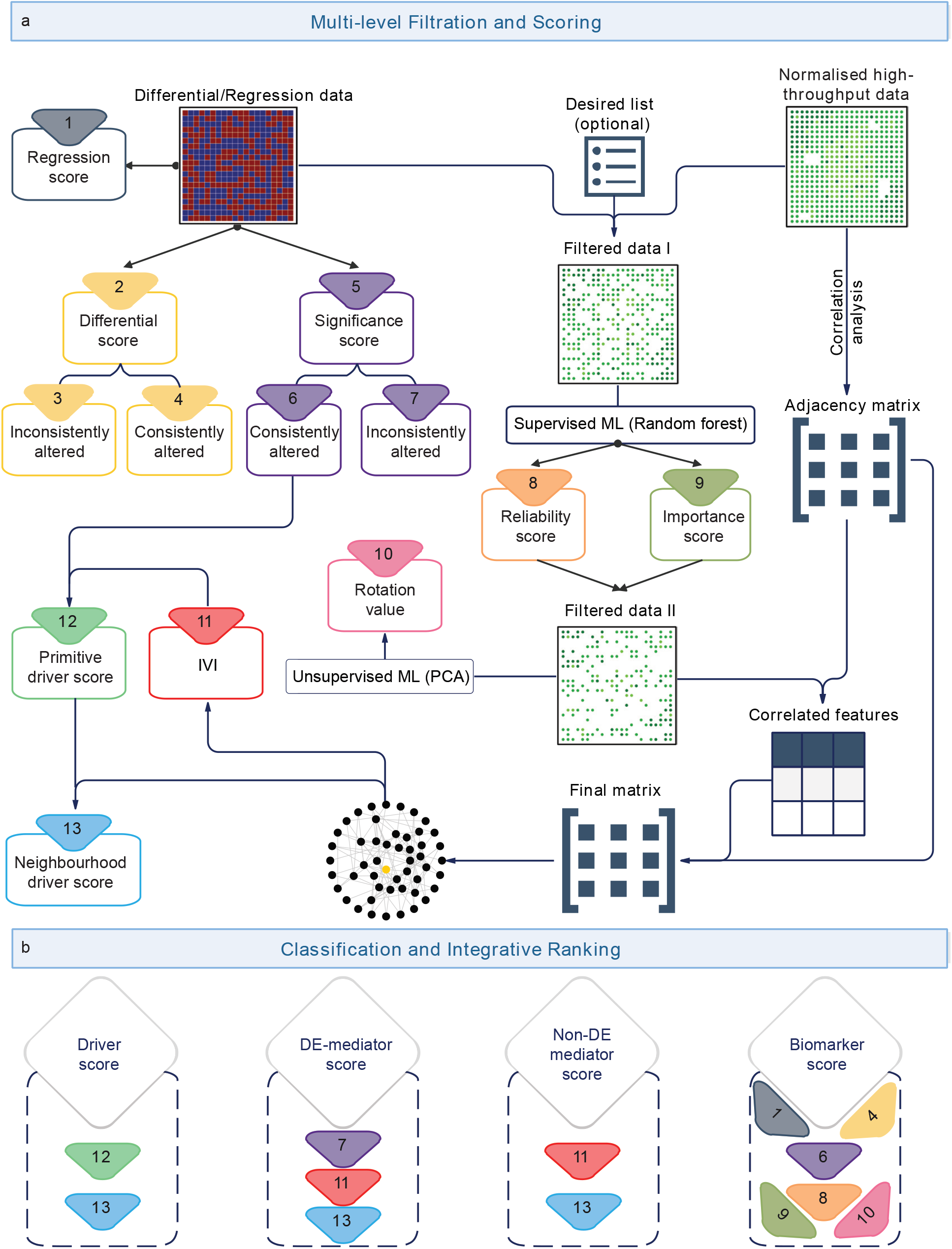
The ExIR model workflow. **a,** ExIR first performs multi-level filtration and scoring of input data (differential/regression data, normalised high-throughput data, and optionally a list of desired features). Differential/regression data are used for generation of multiple scores on the basis of their consistency of alteration between experimental groups to determine primitive and neighbourhood driver scores. Filtered data (I) is subject to supervised ML through the Random Forest algorithm to derive reliability and importance scores, from which additional data filtration is performed (filtered data II). PCA-based unsupervised ML is then performed on this further filtered data to determine a rotation value. Simultaneously, an adjacency matrix is derived from the input normalised data by way of correlation analysis, and a tabulated set of correlated features is derived from intersection analysis of filtered data II. A final association matrix is generated by extending the adjacency matrix by one order using the set of correlated features. A connectivity network is then reconstructed from the final association matrix, which is used to calculate Integrated Values of Influence (IVIs) ^50^ for each feature within the network. IVIs and primitive driver scores are subsequently used to determine final neighbourhood driver scores. **b,** Combinations of scores derived from ExIR’s multi-level filtration and scoring pipeline define feature classifications and ranks. Drivers are classified based on their combined primitive and neighbourhood driver scores; DE mediators (in datasets with >2 groups) are classified based on their IVIs and neighbourhood driver scores and meeting the criteria of having an inconsistently altered significance score; non-DE mediators (not requiring experimental data with >2 groups) do not factor significance scores; and biomarkers are defined by regression scores, consistently altered differential and significance scores, supervised ML-derived reliability and importance scores, and unsupervised ML-derived rotation values.

Unlike other gene/protein prioritisation methods such as Endeavour ^18^ and ToppGene ^19^, which require a seed or training set to derive correlated features, detection of drivers, biomarkers and mediators by ExIR relies only on interrogation of experimental data through both supervised and unsupervised ML approaches (**Fig. 3a, b**). The supervised random forest ML process prioritises genes that would significantly affect inter-group variation between samples. ExIR then reiteratively places more weight on the genes with a greater potential to differentiate all samples from each other, regardless of which group they belong to. Accordingly, those genes that have more sensitivity to inter-group variation and to the severity of the phenotype within each group (*i.e.* intra-group variation) are assigned a higher rank.

We benchmarked ExIR’s performance against other feature prioritisation methods in ranking influential driver and biomarker genes/proteins using a diverse set of previously-published high-throughput transcriptomic and proteomic datasets representing several disease states (glioblastoma ^20^, schizophrenia ^21^, breast cancer ^22,23^, thyroid cancer ^22^, lung adenocarcinoma ^22,24^ and hepatocellular carcinoma ^22^) or physiological processes (oogenesis ^25^). The experimental modalities tested included bulk- and single-cell RNA sequencing, RNA microarray, as well as proteomics datasets, and all processes investigated have at least some known ground-truth drivers and biomarkers (**Extended Data Fig. 2a-d and Supplementary Notes**). For driver and biomarker gene/protein prioritisation, ExIR outperformed all other methods (ToppGene ^19^, Endeavour ^18^, GeneMANIA ^26^ and log fold-change for driver features; mutual information ^27^ [MI], Student’s *t*-test, and point-biserial and Spearman correlation coefficients for biomarker features) in correctly ranking the same set of ground-truth features in all but one dataset analysed (TCGA THCA) as determined by receiver operating characteristic (ROC) analyses (**Extended Data Fig. 2e-h**). In addition to generally outperforming all other feature classification and ranking methods, our application of ExIR to the Cancer Genome Atlas lung adenocarcinoma RNA-sequencing dataset suggested *EMP2* as a potential novel biomarker, which was supported by immunohistochemical data from the Human Protein Atlas database ^28^ (**Extended Data Fig. 3**).

An additional utility of ExIR is its capacity to determine a feature class we refer to as ‘mediators’. Mediators are features that may or may not be differentially expressed/abundant, but have key roles in the propagation of information between nodes corresponding to driver genes in the ExIR-generated network. Thus, mediators are predicted to directly and/or indirectly associate with major drivers of the biological process in question. To our knowledge, only one other model (Machine Learning-Assisted Network Inference, MALANI) ^29^ has so far been described with the capacity to compute mediator features, and has so far only been utilised in the context of breast cancer ^29^. Benchmarking of ExIR’s capacity for mediator feature detection against MALANI by overrepresentation analysis using the TCGA breast cancer RNA-sequencing dataset demonstrated that ExIR significantly outperformed MALANI in identification of mediators associated with breast cancer-related GO biological processes (GO-BPs) and KEGG pathways. Extension of this analysis to mediator identification in other disease high-throughput datasets identified, on average, that >30% of mediators corresponded to disease-relevant GO-BPs and KEGG pathways (**Extended Data Fig. 2i-l, Extended Data Tables 1-2**). Taken together, our benchmarking demonstrates ExIR’s powerful capacity to infer and rank driver, biomarker and mediator features from high-throughput datasets without reliance on external sources of information.

### Implementation of the ExIR workflow to systems-level analyses of the sgsh zebrafish CNS highlights diverse functional contributors to MPS IIIA pathology

In order to prioritise differentially-expressed/abundant features resulting from RNA-seq and quantitative proteomics analyses of young wild type and *sgsh* brain in a manner independent of external ontological data, we applied the ExIR workflow to each dataset and the time points therein. Ranks were thus called for upregulated/accelerating (**Extended Data Fig. 4a**) and downregulated/decelerating genes (**Extended Data Fig. 4b**) and proteins (**Extended Data Fig. 4c, d**). The top three accelerating transcriptional drivers of the MPS IIIA disease state in young *sgsh* brain *(i.e.* upregulated features that have the highest driving potential within the generated ExIR network, and are surrounded in that network by other prioritised driver features) were *baiap2b; ppargc1b,* and *tet1* (**Extended Data Fig. 4a**). The top three ranked accelerating transcriptional biomarkers (*i.e.* upregulated genes that exhibit significantly and consistently altered behaviour in different conditions, and have the greatest potential to discriminate between conditions) somewhat overlapped with the corresponding accelerating driver population; these were *tyrp1b, baiap2b*, and *tet1*. Accordingly, the top three decelerating transcriptional drivers in the young *sgsh* brain were the IEGs *junbb* and *npas4a*, and the serine-threonine kinase *sik1*. Top decelerating biomarkers in the young *sgsh* brain included the predicted long non-coding RNA LOC100535167, as well as the IEGs *npas4a* and *junbb* (**Extended Data Fig. 4b**).

In the aged *sgsh* brain, the top accelerating transcriptional drivers were *pmela*, *tspan36* and *acvr2aa* – all pigmentation/melanocyte-associated genes ^30–32^ – and top accelerating transcriptional biomarkers included *pmelb*, *pmela* and *pdgfrl* (**Extended Data Fig. 4e**). *In situ* hybridisation demonstrated no detectable expression of *pmela* in the wild type zebrafish brain, but strong expression specifically within the choroid plexus in *sgsh* brain (**Extended Data Fig. 1e-f**). Top aged decelerating transcriptional drivers included the predicted ATP-binding protein *si:ch73-236c18.5,* a collagen alpha-1 (XXVI) chain-like protein LOC101885935 and the kinesin family member *kif21a* involved in microtubule-dependent intracellular transport and regulation of the axonal cytoskeleton ^33^. ExIR-prioritised downregulated transcriptional biomarkers in aged *sgsh* brains included the endosome-associated *rab22a*, *si:ch73-236c18.5*, and the ubiquitin ligase *fbxo45* that regulates proteasomal degradation at the synapse ^34^ (**Extended Data Fig. 4f**).

ExIR was additionally used to rank and classify the differentially abundant proteins identified between both young and aged wild type and *sgsh* brains. The top *sgsh*-enriched driver and/or biomarker proteins in young brains were Gnsa, Cpvl, Ctsla, and Dnase2 (**Extended Data Fig. 4c**); while the top wild type-enriched protein drivers and/or biomarkers were Arsa, Gaa2, Mdkb and Mfap4.2 (**Extended Data Fig. 4d**). While the *sgsh*-enriched proteins in the aged brain largely overlapped with those observed in the young brain, the ExIR rankings of these proteins was noticeably altered; top *sgsh*-enriched driver and/or biomarker proteins in aged brain were Hexb, Napsa, Ctsba, Dnase2, Stm and Naga (**Extended Data Fig. 4g**). Top WT-enriched driver/biomarker proteins in the aged brain included Arsa, Palm1a, Mdkb and Ckmt2a (**Extended Data Fig. 4h**).

In addition to classification and ranking of driver and biomarker features, the ExIR model infers a class of genes – mediators – which represent nodes in the generated network important for the flow of information between prioritised drivers (**Fig. 3b**). 365 transcriptional mediators were present in both young and aged RNA-sequencing datasets. We performed STRING analysis on these genes and, due to their large number and broad functional diversity, applied stringent filters for interaction score. Following exclusion of singletons from the STRING network, 34 conserved mediators remained (**Extended Data Fig. 5a**). Gene ontology overrepresentation analysis showed that these genes were functionally enriched in mitophagy, nucleophagy and microautophagy, vesicle fusion at the presynaptic active zone membrane and tetrahydrofolate metabolism (**Extended Data Fig. 5b**). Application of ExIR to the proteomic analysis of *sgsh* CNS yielded a large number of protein mediators between young and aged timepoints (620 shared out of 1777 young and 1689 aged mediators (**Extended Data Fig. 5c**). The resulting STRING network assembled from cross-timepoint proteomic mediators, comprising 99 nodes excluding singletons, exhibited a far greater degree of interactivity (**Extended Data Fig. 5c**).Only one protein was represented in the top 10 mediators of both time points *(bloc1s5,* rank #8 young, #9 aged) (**Extended Data Fig. 5d-e**). Major mediator functional clusters represented proteins involved in ribosomal assembly and translation initiation, vesicle fusion and exocytosis (including of synaptic vesicles), nucleotide/nucleoside biosynthesis, and amino acid biosynthesis (**Extended Data Fig. 5f**). By way of the definition of a mediator in ExIR, none of the genes or proteins in the above data are differentially expressed or abundant due to the low dimensionality of these datasets (**Fig. 3a, b**) – thus, it is notable that despite the ExIR network being assembled purely on the basis of normalised expression/abundance values, many biological processes associated with MPS IIIA pathology are inferred by ExIR without input from extraneous knowledge sources.

### Immediate early gene perturbation indicates deficient synaptic activity underlies progression of MPS IIIA neuropathology

With the assistance of feature refinement and prioritisation by ExIR, our transcriptomic analyses highlight profound dysregulation of factors involved in synaptic neurotransmission in the *sgsh* brain, as well as broadly downregulated expression of neural activity-related IEGs (**Fig. 1a-c, f-i’**). Constitutively reduced IEG expression across the *sgsh* CNS is a logical consequence of reduced synaptic activity. Examination of IEG expression in wild type brain by *in situ* hybridisation against *egr1* transcripts demonstrated particularly strong expression in the midbrain and hindbrain (**Fig. 1f-g’**), which was largely ablated in the *sgsh* mutant brain (**Fig. 1h-i’**). We thus sought to study synapses at the ultrastructural level in these regions in adult wild type and *sgsh* brains using TEM, and observed that wild type axon terminals generally exhibited abundant synaptic vesicles (SVs) in both the active zone and reserve pool (**Fig. 4a**). Contrastingly, axon terminals in *sgsh* neurons frequently exhibited greatly reduced numbers of reserve pool SVs (RP-SVs), with most observed SVs located at the active zone(s) (**Fig. 4b, c**). This, in combination with the previously-observed accumulation of dense axonal microtubule networks (**Fig. 2n, n’**), suggests that RP-SV depletion in the *sgsh* brain may be associated with impaired SV trafficking. To elaborate on this, we quantified the number of cerebellar Synapsin-1/2 immunoreactive glomeruli (tripartite synapses of presynaptic axon terminals of mossy fibres and Golgi cells, and postsynaptic granule cells) in adult wild type and *sgsh* zebrafish (**Fig. 4d-e**). Synapsins are major components regulating the organisation of SV clusters that occur at presynaptic terminals ^35^; and perturbed synapsin expression or function impairs SV availability at the reserve pool ^36,37^ *sgsh* cerebelli had significantly fewer Syn1/2^+^ glomeruli throughout the granule cell layer compared to wild type siblings (**Fig. 4f**). Taken together, downregulation of multiple IEGs in the *sgsh* CNS coincides with reduced synapsin immunoreactivity, which collectively manifests as a reduction in RP-SVs in the *sgsh* mutant CNS. Examination of basal expression levels of several IEGs (i.e. in the context of normal levels of neural activity) by RT-qPCR (**Extended data Fig. 6a**) and *in situ* hybridisation (**Extended data Fig. 6b-e**) indicated they were not differentially expressed in larval *sgsh* brains as they are in the adult CNS. This demonstrates that overtly deficient IEG expression in the adult *sgsh* brain is a result of a progressive phenotype. Despite this, we sought to establish whether abnormalities in neuronal activity could be observed earlier than the later-detected molecular and ultrastructural changes in the adult *sgsh* brain. To dynamically detect early changes in neural activity, we employed live imaging of calcium signalling dynamics using pan-neuronally-expressed H2B-GCaMP6s ^38^ in *sgsh* larvae and wild type siblings acutely administered a high dose of the convulsant pentylenetetrazole (PTZ) in conjunction with rapid, intermittent (every 0.5s) photic stimulation (**Fig. 4g**). This was performed to challenge the neural circuitry of the larval *sgsh* brain, and induce stereotypic patterns of neuronal firing. Like other mechanisms of ectopic neural stimulation – both electrical ^39^ and pharmacological ^40^ – PTZ is able to induce mobilisation of RP-SVs beyond what is observed in physiological behaviours ^41^. Treatment of wild type and *sgsh* larvae with PTZ resulted in stronger *npas4a* expression in wild type larval brains than in *sgsh* brains (**Extended Data Fig. 6f-h**) suggesting a difference in neuronal excitability or capacity for full depolarisation. GCaMP imaging demonstrated that PTZ-treated wild type larvae generally exhibited a single, large brain-wide depolarisation event, followed by an extended refractory period in which GCaMP6s fluorescence was heavily reduced (mean seizure count = 2.083 ± 0.3128 SEM) (**Fig. 4h, j**). In contrast, PTZ-treated *sgsh* larvae exhibited multiple low-grade seizures (mean seizure count = 4.333 ± 0.3404 SEM) with short refractory periods between each seizure (**Fig. 4i, j**). This observation further supports the hypothesis that synaptic activity exhibits underlying susceptibility to dysfunction in the developing *sgsh* CNS. Collectively, these data indicate that the *sgsh* zebrafish exhibits a susceptibility to perturbed neuronal activity well-preceding the onset of later IEG dysregulation and functional neurodegeneration.

**Fig. 4.**
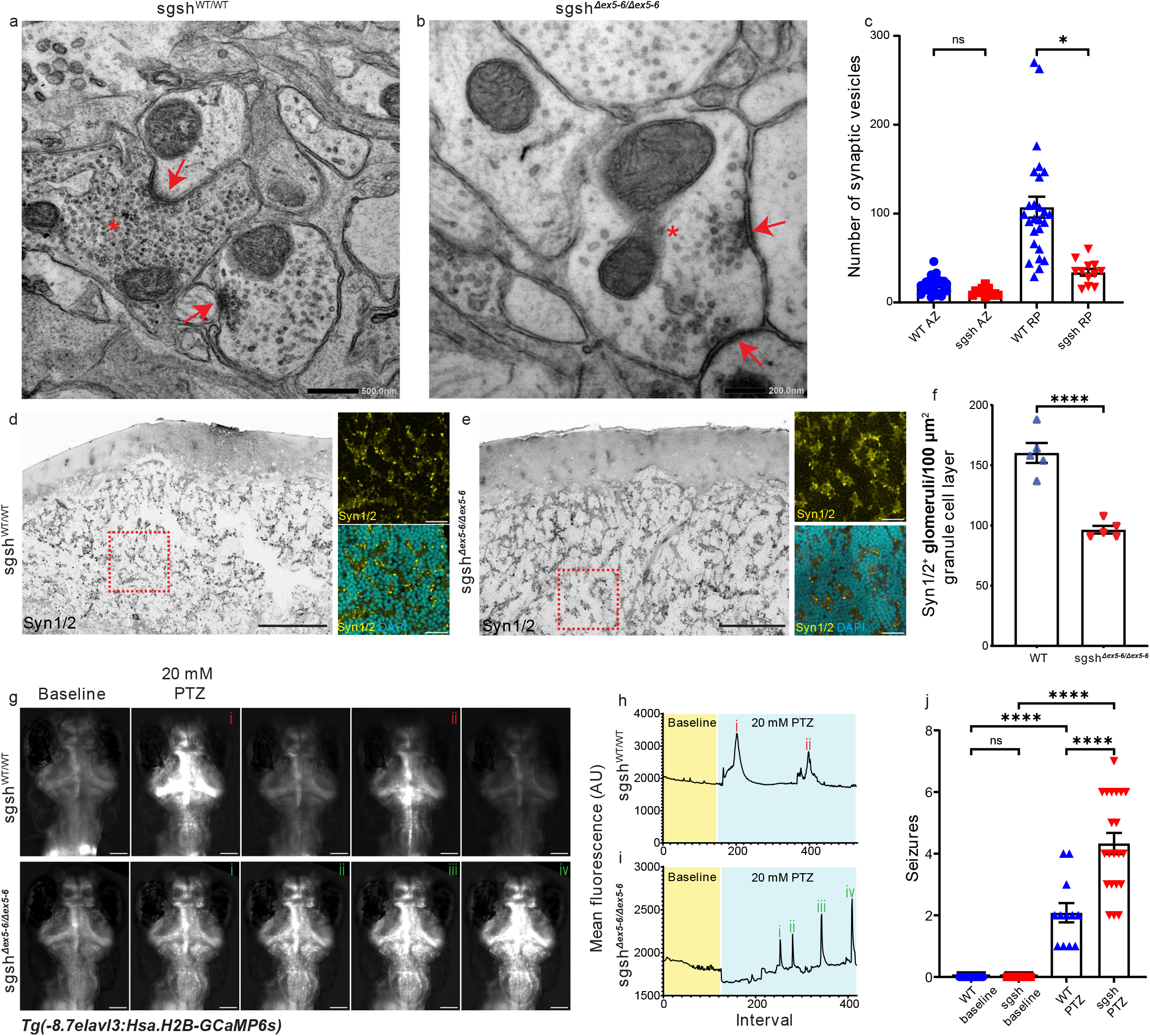
Structural and functional aberrations in *sgsh* synapses result from impaired synaptic vesicle localisation to the synapse and maintenance of the synaptic vesicle reserve pool. **a,** TEM of axon terminal (red asterisk) and synapses (red arrows) in wild type brain. Scale bar 500 nm. **b,** TEM of axon terminal (red asterisk) and synapses (red arrows) in *sgsh* brain. Scale bar 200 nm. **c,** Quantification of synaptic vesicles in the synaptic active zone (AZ) and reserve pool (RP) of wild type and *sgsh* axon terminals. ns = not significant, * *P* = 0.0325; Kruskal-Wallis test with Dunn’s multiple comparisons test. **d,** Synapsin-1/2 immunoreactivity in the wild type cerebellar granule cell layer; scale bar 100 μm. Panels are high-magnification images of boxed region showing Syn-1/2^+^ cerebellar glomeruli; scale bars 10 μm. **e,** Synapsin-1/2 immunoreactivity in the *sgsh* cerebellar granule cell layer; scale bar 100 μm. Panels are high-magnification images of boxed region showing Syn1/2^+^ cerebellar glomeruli; scale bars 10 μm. **f,** Quantification of Syn1/2^+^ glomeruli in 100 μm^2^ of wild type and *sgsh* granule cell layer. Each data point represents one animal; **** *P*< 0.0001, Student’s *T*-test. **g,** GCaMP6s fluorescence in 7 dpf *Tg*(*−8.7elavl3:H2B-GCaMP6s)* brain at baseline and after 20 mM PTZ administration in wild type (top) and *sgsh* homozygotes (bottom). Scale bars 100 μm. **h,** Representative whole-brain GCaMP6s fluorescence trace in a wild type 7 dpf larva at baseline and following 20 mM PTZ administration. Red (i) and (ii) are seizure peaks in **g**, top panels. **i,** Representative whole-brain GCaMP6s fluorescence trace in a homozygous *sgsh* 7 dpf larva at baseline and following 20 mM PTZ administration. Green (i) and (ii) are seizure peaks in **g**, bottom panels. **j,** Quantification of seizures in wild type and *sgsh* homozygous larvae after PTZ administration. ns = not significant; **** *P* < 0.0001; Ordinary one-way ANOVA and Šídák’s multiple comparisons test with a single pooled variance.

## Discussion

The pathological origin of mucopolysaccharidoses – that is, impaired GAG catabolism (HS in the case of MPS IIIA) – has long been known. Despite this, it remains unclear how this manifests as the characteristic progressive functional neurodegeneration observed in MPS patients. Several studies have implicated multiple factors contributing to disease progression, including microgliosis and neuroinflammation ^1,8,42^, CNS atrophy ^43^, and more recently an increasing recognition of the role of functional synaptic impairments ^44–46^. However, most studies have looked at these features largely in isolation; here, we have performed the first transcriptomic and proteomic characterisation of the CNS of an animal model of MPS IIIA, the *sgsh* zebrafish ^8^, with the aim of globally determining the features associated with disease progression and defining the diverse cellular and molecular perturbations that accumulate in the MPS IIIA brain. The *sgsh* zebrafish brain exhibits profound accumulation of LAMP1-immunoreactive organelles as well as abnormal lipid storage in major neuronal populations, reflecting the increased endolysosomal burden that results from impaired HS catabolism without negative regulation of its biosynthesis. This observation was supported by our quantitative proteomics analyses, which highlighted massive enrichment of proteins involved in lysosomal substrate degradation, indicative of the progressive accumulation of lysosomes and lysosome-like organelles. Such a phenotype would be expected to correlate with formation of dense microtubule networks, which are required for endocytosis and lysosome formation; such networks were widely observed upon ultrastructural examination of myelinated axons in the *sgsh* brain. A consequence of this relates to the requirement of microtubule-dependent intracellular trafficking of neurotransmitter-containing vesicles ^47^; as endocytosis and lysosomal sequestration of extracellular HS in MPS IIIA requires increased microtubule-based cytoskeletal remodelling, this phenotype may interfere with intracellular SV trafficking and directly relate to the observed reduction in SV localisation at presynaptic terminals. This is supported by reduced synapsin-1/2 immunoreactivity observed in neuronal populations in the *sgsh* brain, and a consequent significant reduction in the number of SVs present in the presynaptic reserve pool. Synaptic perturbation in the *sgsh* brain was further evidenced in our transcriptomic analyses, wherein a profound relative brain-wide reduction IEG expression (a transcriptional hallmark of postsynaptic activation ^10,48^) was observed in both young and aged *sgsh* brains. While expression levels of IEGs are expected to fluctuate depending on stimulatory input, constitutively diminished expression in homozygous *sgsh* brains compared to wild type siblings indicates the presence of fundamental perturbations in stimulus-response and excitation-transcription coupling. A recent study has linked lysosomal disruption to impaired synaptic activity in a mouse model of MPS IIIA ^45^, but ascribed this to presynaptic accumulation of alpha-synuclein and the DNA J-protein CSPα. However, these features do not explain the strongly conserved synaptic phenotype between the MPS IIIA mouse model and our *sgsh* zebrafish, as zebrafish lack an orthologue for alpha-synuclein ^49^ and the zebrafish orthologues for mouse CSPα*(dnajc5aa, dnajc5ab)* were neither differentially expressed at the transcript level nor differentially abundant at the protein level in the *sgsh* zebrafish brain. Though this may be explained by variability in neurobiology between the two species, it is unlikely that divergent pathological mechanisms would lead to the same resulting phenotype. This suggests that alternative mechanisms, such as the endolysosomal impairment of SV trafficking, are more likely conserved pathways involved in driving MPS IIIA progression in both model systems as well as in patients.

Given the lack of a concrete explanation underpinning the pathobiology of MPS IIIA progression in the CNS, we sought an unbiased manner by which to determine putative key features from our ‘omics data that drive the disease state. To this end, we developed a novel bioinformatic tool, ExIR (Experimental data-based Integrate Ranking); ExIR is a versatile model that simultaneously extracts, classifies, and prioritizes candidate features (*e.g.* genes, proteins) from high-throughput experimental data independent of external, curated knowledge sources. ExIR initially performs several filtrations of input normalised data to derive several different ranking scores, which are then used in multiple combinations to generate summary scores for feature classification and ranking as drivers, biomarkers and mediators of biological processes. From a biological perspective, manipulation of driver features prioritized by ExIR (potentially alongside co-manipulation of associated mediators) could have the most prominent impact on the progression of a biological process/disease as well as phenotype manifestation. While highly-ranked ExIR biomarkers are anticipated to have the highest sensitivity to a biological condition and the severity of the phenotype, their manipulation may not necessarily affect the progression of the process in the way highly-ranked driver features might.

Our comparative analyses show that ExIR robustly outperforms current tools and algorithms in the unbiased prioritisation of ground truth driver, biomarker, and mediator features. The underpinning reasons for the superior accuracy and robustness of our model include (1) co-implementation of both supervised and unsupervised ML techniques; (2) integration of both ML and network-based models; (3) optimisation of mathematical operations for score integration; and (4) independence from potentially-confounding external sources of information. Overall, ExIR has the potential to provide significant momentum to biological discoveries through streamlining classification and prioritisation of features in high-throughput data and allowing for rapid generation of hypothesis from refined and more interpretable data. As it is a standalone computational model, we expect ExIR to be of broad and multidisciplinary utility, for example as an efficient tool for in guiding precision medicine as well as for the expedition of diagnosis and treatment of rare diseases through specific and sensitive ranking of biomarkers and drug targets.

Taken together, our work utilises a cross-disciplinary approach to develop from multiple sources of high-throughput data a unified hypothesis integrating the diverse features underlying the origins and progression of neurological decline in MPS IIIA. As the mechanism for MPS IIIA disease progression we hypothesise does not exhibit any features unique to this particular disease beyond the requirement for massive endolysosomal storage of heparan sulfate, this mechanism may be generalisable to a broad range of neuronopathic/synaptopathic storage disorders and thus may have far-reaching implications for the development of much-needed therapies for this class of disease.

## Methods

### Development of the ExIR model

ExIR is comprised of two major components. The first is the multi-level filtration and scoring which calculates several scores directly from the supplied experimental data. Normalised experimental data is filtered, and both supervised and unsupervised machine-learning (ML) algorithms are employed to extract and assign weight to the prominent biological features of the supplied dataset (*e.g.* transcriptomics, proteomics). Additionally, network techniques are employed for assessing the importance of association between genes within the constructed network. The second component is the classification and integrative ranking, where biological features are classified according to the integration of different combinations of scores calculated in the first part of the model.

#### Mathematical basis of score integration

As demonstrated previously ^50^, basic arithmetic operators can be employed for the integration of different scores. *Addition* is appropriate for the combination of two co-essential (*i.e.* associated) measurements with the purpose of compensating for deficiencies in individual scores. *Multiplication* is appropriate for the integration of two scores provided that the objective is to synergise their effects ^50^, and has been used by other gene prioritisation methods such as MetaRanker ^51^ to compute the final rank for each gene in a list. Accordingly, we employed these operators for the integration of different scores. All scores were range-normalised while maintaining their relative weight ratio using the Min-Max gene scaling algorithm ^52^.

#### Differential/regression data scoring

The differential score in calculated by applying the *addition* function to the absolute of all differentiation values:

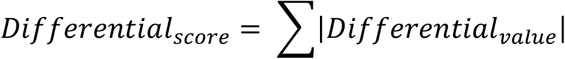

The *Differential_value_* is the user-calculated differential expression/abundance of genes/proteins. For example, *Differential_value_* may be log2FC corresponding to DEGs in a transcriptomic assay. If the study is a time-course experiment, or includes >2 step-wise conditions, it is possible to include all sets of differential values corresponding to different pairwise differential analyses. Similarly, the regression score is calculated as the sum of all regression values:

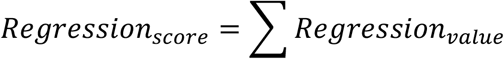

where *Regression_value_* is the previously user-calculated R^2^ value of genes. Importantly, the R^2^ value is calculated in regression analyses in terms of the target variable, which in biological contexts corresponds to the expression/abundance level of the gene/protein in question. The explanatory variable must be determined according to the specific biological context of the data in question. In a time-course study, for example examination of developmental progression in an organism, time is considered the explanatory variable used for prediction of the amount of the dependent variable *(e.g.* gene expression level). However, this score applies only to time-course and multi-condition datasets and is thus optional for the development and execution of ExIR.

The significance score is calculated by combining the log-transformed statistical significance (*e.g.* P-value, adjusted P-value, false discovery rate) of differential (and/or regression) values as follows:

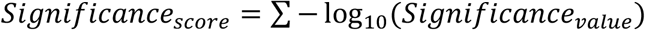

where *Significance_value_* refers to statistical significance values prior calculated by the user corresponding to the assay type in question.

#### Normalisation and filtration of experimental data

Experimental data input into ExIR should first be normalised according to the nature of the experiment, to account for probable biological and technical sources of experimental variation. Unless otherwise specified, no further normalisation is performed in the ExIR workflow. If input experimental data is non-normalised, ExIR will optionally perform a log-transformation to normalise said data. However, as not all data would benefit from a log-transformation, we recommend prior normalisation of the data before implementation of the ExIR workflow. Following this, experimental data is filtered by either DEGs or, if provided, a list of desired genes.

#### Supervised ML

The random forest algorithm ^53^ was selected for implementation in the ExIR workflow due to its appropriateness for problems of large *p* (*i.e.* number of genes) and small *n* (*i.e.* number of variables/samples), as well as its variable importance measures which are useful for gene ranking ^54^. ExIR builds a random forest model from supplied experimental data using the ranger R package ^55^, with (by default) the number of trees set to 10,000 and the *mtry* parameter (*i.e.* the number of variables to possibly split at each node) set to the square root of the number of genes in the filtered experimental data. To calculate the importance of gene scores, we applied the impurity-corrected mode; an unbiased, fast and memory-efficient method for the evaluation of the importance of genes in a random forest classification model ^56^. In combination with other methods, impurity correction can be used to estimate statistical significance of ExIR-derived importance scores. Accordingly, P-values of importance scores are computed using the permutation-based method proposed by Altmann *et al.*^57^, with the number of permutations by default set to n=100. Finally, experimental data is further filtered according to the significant genes output from the random forest classification. All parameters described above are user-adjustable as required.

#### Unsupervised ML

In parallel to the supervised ML, a correlation analysis is performed on the unfiltered experimental data using the Spearman algorithm in order to investigate the association of genes with each other in a systematic manner. In ExIR, genes are first ranked using base R functions. Subsequently, the correlations between all pairs of genes are calculated using the coop R package (https://cran.rproject.org/package=coop), which enables the vectorisation of operations and usage of multiple cores whenever possible. This consequently increases the speed of correlation analyses by several orders of magnitude. Correlation data is assembled in an adjacency matrix, which is then refined against the genes provided in the filtered experimental data to extract only positive correlations between these genes and all other genes within the dataset. From this, a list of correlated genes is thus obtained. Similarly, the initial unfiltered adjacency matrix is filtered against the list of correlated genes, and an ultimate adjacency matrix is generated comprising first- and second-order correlated genes from the filtered experimental data. Such a step-wise assessment of correlation is employed to increase the accuracy and speed, while reducing the computational intensity, of subsequent network reconstruction and analyses.

#### Network reconstruction and calculation of Integrated Values of Influence (IVIs)

The igraph R package ^58^ was used to reconstruct the association network from the ultimate adjacency matrix derived by correlation analysis. IVIs of all nodes (*e.g.* genes, proteins) within the network were computed using the influential R package (https://cran.r-project.org/package=influential)^50^.

#### Calculation of primitive and neighbourhood driver scores

The primitive driver score is calculated by synergising the effects of IVI and the significance score of consistently-altered genes using the multiplication operator:

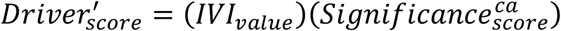

where 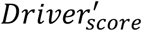 represents the primitive driver score and 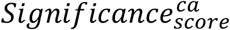 represents the consistently altered significance score. Subsequently, the primitive driver scores are mapped onto the network nodes. Accordingly, the neighbourhood driver score of a given node *i* is calculated by computing the additive product of primitive driver scores of all first-order connectors to that node:

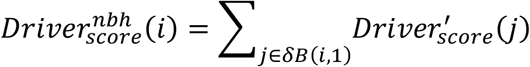

The 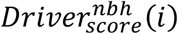 represents the neighbourhood driver score of node *i*; while *δB*(*i*, 1) is the set of nodes with distance 1 from node *i*. 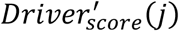 is the primitive driver score of node *j*, a first-order connector node to node *i*.

#### Gene classification and scoring

From the scores calculated in the first part of the ExIR workflow, four separate classes of genes are defined based on variable integration of different score combinations. The final driver score is calculated by synergising the effects of primitive and neighbourhood driver scores under the assumption that the top candidates would be surrounded in the network by high-ranked neighbours ^59^:

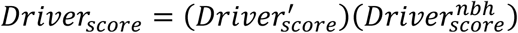

Driver genes are themselves divided into two subtypes based on the directionality of their effects; *accelerator* drivers are up-regulated in a biological process, while *decelerator* drivers are down-regulated.

The biomarker score is achieved by applying the multiplication operator on the differential, regression, statistical significance and ML-derived scores:

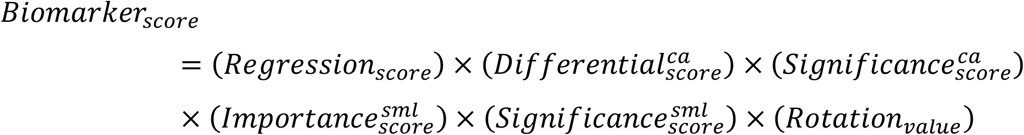

Here:

- 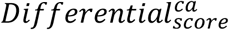 is the *Differential_score_* of consistently altered differentially expressed/abundant features;
- 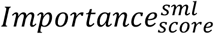 and 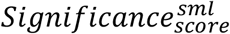, refer to supervised ML-derived importance and significance scores, respectively;
- *Rotation_value_* represents the rotation value derived from unsupervised ML.

Similar to drivers, biomarkers are separately classified as up- or down-regulated.

Mediators are classified as either DE or non-DE. DE mediators are those that are DE but in an inconsistent manner; accordingly, the DE mediator score is calculated by integrating the significance scores of inconsistently altered DE features with their IVIs and neighbourhood driver scores:

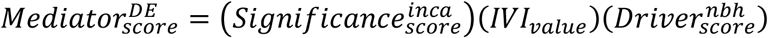

where the 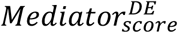 represents the DE mediator score and 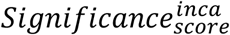 represents the significance score of inconsistently altered DE features. In contrast, non-DE mediator are those that are not DE at all *(i.e.* in any experimental group), and thus this score is calculated by synergising the effects of the IVI and neighbourhood driver score:

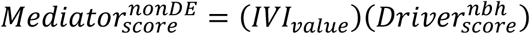

#### Calculation of the statistical significance of prioritised genes

Scaling and normalisation of scores was achieved by standardisation using the Z-score transformation method ^60^ through base R functions. Next, P-values are calculated based on the computation of Z-score probability distributions, and adjusted using the Benjamini and Hochberg algorithm via the stats R package.

### Data preparation for ExIR evaluation and benchmarking

For evaluation and benchmarking of ExIR, the following criteria were required for selection of datasets:

1. The dataset should correspond to a disease/biological process with >40 curated driver genes in the DisGeNET database or Gene Ontology resource;
2. The dataset should have >100 samples/cells;
3. The dataset should include at least 2 conditions (*e.g.* diseased vs. unaffected, different time points);
4. The dataset should have a comparable number of samples/cells within each condition.

Accordingly, the following datasets were used to evaluate ExIR and demonstrate its applicability in the extraction, classification and prioritisation of genes from microarray, bulk and single-cell RNA-seq (scRNA-seq) data.

#### Glioblastoma (GBM) dataset

This is a scRNA-seq dataset generated using plate-based protocols from four patients with confirmed cases of primary GBM and comprises 3,589 cells ^20^. Tissue samples for this dataset originated from either the tumour core or the peritumoural cortical space. Major classes of cells, including neoplastic and non-neoplastic cells, were identified using immunopanning ^61^ and further confirmed by comparison with other single-cell and bulk RNA-seq data. The dataset was originally normalised based on the read counts generated by HTSeq ^62^ and filtration of genes with very low counts. Amongst all cell types previously identified, we selected two major subsets of cells including periphery regular (non-neoplastic) cells (n=1,184 cells) as the ‘normal’ set and tumour neoplastic cells (n=1,029 cells) as the cancer set, and filtered out genes with zero counts across all selected cells for downstream analysis.

#### Oogenesis dataset

This is a scRNA-seq dataset generated using plate-based protocols from fetal mouse ovaries at three developmental stages (E12.5, E14.5 and E16.5), together encompassing 19,144 FACS-sorted high-quality murine female germ cells ^25^. The dataset was originally normalised in Seurat ^63^, as was DEG detection between all time-points.

#### TCGA BRCA

This is a bulk RNA-seq of breast cancer generated by the TCGA project. Here, only primary tumour (#1,095) and solid tissue normal (#113) samples were retrieved using the TCGAbiolinks R package ^64^. The raw RNA-seq data were pre-processed based on the Array-Array Intensity Correlation (AAIC) method in TCGAbiolinks with default parameters (r>0.6). Processed data underwent quantile normalisation using the default parameters of TCGAbiolinks function for downstream analyses. All other TCGA datasets used in this work were pre-processed and normalised according to the same methods and parameters applied to this dataset.

#### TCGA THCA

This is a bulk RNA-seq dataset of thyroid carcinoma generated by the TCGA project. Samples retrieved were primary tumour (#505) and solid tissue normal (#59).

#### TCGA LUAD

This is a bulk RNA-seq dataset of lung adenocarcinoma generated by the TCGA project. Samples retrieved were primary tumour (#515) and solid tissue normal (#59).

#### TCGA LIHC

This is a bulk RNA-seq dataset of liver hepatocellular carcinoma generated by the TCGA project. Samples retrieved were primary tumour (#371) and solid tissue normal (#50).

#### Schizophrenia dataset

This is a total RNA microarray dataset generated from dysfunctional dorsolateral prefrontal cortex layer 3 parvalbumin neurons in 36 matched pairs of schizophrenia and unaffected cases using the Affymetrix Human Genome U219 array ^21^. This dataset contains 141 samples including 71 healthy and 70 schizophrenia samples. The raw microarray data were retrieved from the GEO database (GSE93577) utilising the GEOquery R package ^65^ and log-transformed prior to downstream analyses.

#### LUAD proteomics dataset

This is a proteomics dataset generated from primary LUAD samples with paired non-cancerous adjacent tissues from treatment-naive patients by means of high-performance liquid chromatography-mass spectrometry (HPLC-MS) and label-free quantification ^24^. This dataset contains 206 samples including 103 normal and 103 LUAD samples. The MaxQuant-based pre-processed data was retrieved from the Integrated Proteome Resources (IPX0001804001).

#### BRCA proteomics dataset

This is a proteomics dataset (PXD002057) generated from human breast cancer cell lines SKBR3 and BT474 and their lapatinib-resistant derivative cells by means of nano-scale HPLC-MS and label-free quantification ^23^. This dataset contains 20 samples including 10 benign and 10 malignant samples. The MaxQuant-based pre-processed data of this dataset was retrieved from the LFQ-Analyst website (https://bioinformatics.erc.monash.edu/apps/LFQ-Analyst/).

### Differential expression/abundance analyses in ExIR benchmarking

#### GBM dataset

Differential expression analysis (DEA) was performed using the DEsingle R package ^66^. DEsingle uses a zero-inflated negative binomial model to estimate the proportion of dropout and real zeroes in order to accurately identify DEGs. Subsequently, the normalised fold change values were log_2_-transformed to identify up- and down-regulated genes. DEGs with Padj > 0.05 were filtered out.

#### TCGA datasets

All TCGA datasets used in this work underwent DEA using the TCGAbiolinks R package, which implements functions of edgeR ^67^. Specifically, a common negative binomial dispersion was first estimated across all genes, and a negative binomial log-linear model was fit to the read counts for each gene. Then, pair-wise tests for differential expression between the two groups were performed. All P-values were adjusted, and DEGs with Padj > 0.05 were filtered out.

#### Schizophrenia dataset

Microarray DEA was performed using the limma R package ^68^, where a linear model was first fit to the expression data of each probe, followed by computing the contrasts of the fitted models with moderated empirical Bayes statistics. DEGs with Padj > 0.1 were filtered out.

#### Proteomics datasets

The proteomics data were analysed using the R package DEP ^69^. More precisely, initially each dataset was filtered for proteins that had a maximum of 20 percent missing values in at least one condition within each dataset. Next, the variance of each dataset was normalized followed by a missing value imputation using the “man” algorithm. Lastly, the differential abundance of proteins was calculated using the limma method, P-values were adjusted using the Benjamini and Hochberg algorithm. All P-values were adjusted, and differentially abundant proteins with Padj > 0.05 were filtered out. In the case of LUAD proteomics dataset the differentially abundant proteins with |log_2_FC| < 1 were filtered out to maintain the most prominent characteristics of the disease for downstream analyses and benchmarking.

### Evaluation of driver prioritisation in ExIR

To assess the performance of ExIR in driver gene prioritisation, the sets of curated driver genes of the above datasets were retrieved from either DisGeNET v7 ^70^ or the Gene Ontology resource ^71^ and considered as the ground truth (**Extended Data Table 3**). Additionally, the ground truth drivers of the TCGA datasets were complemented with the driver genes proposed by the MutPanning web server (http://www.cancer-genes.org/)^72^. Next, an intersection analysis was performed to identify common genes between the sets of ground truth driver genes and the sets of significantly up-regulated genes in the selected datasets. As genes with the most statistically significant differential expression are more likely to be driver genes ^73^, a set of DEGs with the least significant Padj and with the same length as their respective true positive set were selected as true negatives. Then, these sets of true positive and negative genes were combined an input into ExIR as the desired lists of genes. The outputs were compared with four commonly used driver gene prioritisation methods; log2 fold change (log2FC), GeneMANIA ^26^, Endeavour ^18^ and ToppGene ^19^. The evaluation and comparison of driver gene prioritisation methods were performed based on the receiver operating characteristic (ROC) analyses using the plotROC R package ^74^. The varied threshold in all ROC analyses in this study is the threshold of prioritisation rankings used for defining true positive and negative features via the variation of which a ROC plot is generated for each benchmarked method. This threshold is calculated by default according to the ROC algorithm. To obtain the prioritised driver genes, the default parameters were used to run the GeneMANIA and ToppGene models. For Endeavour, the gene ontologies, Reactome pathways ^75^ and STRING PPIs ^15^ were selected for building the models. Moreover, the training sets required to run the Endeavour and ToppGene models included all genes previously retrieved from the DisGeNET, MutPanning and Gene Ontology databases, except for those genes selected for testing the models (**Figure 2a-b**).

### Evaluation of biomarker prioritisation

To evaluate the performance of ExIR in the sensitive and specific identification of biomarkers, lists of biomarkers for cancer and non-cancer diseases were obtained from the NCI EDRN (https://edrn.nci.nih.gov/biomarkers) and a knowledge-based database of disease-related biomarkers ^76^ (accessed August 19, 2020), respectively. The EDRN has proposed >20 biomarkers for two out of the five cancer types corresponding to the above TCGA datasets – specifically, lung and breast cancer. The lists of protein/proteomic biomarkers were retrieved from the EDRN (**Extended Data Table 4**). Additionally, a list of schizophrenia biomarkers was derived from the database of disease-related biomarkers ^76^ **(Extended Data Table 4)**, which employs a knowledge-driven text-mining approach to extract the biomarkers of a wide variety of diseases. The common features between these sets and the previously obtained lists of significantly up-regulated genes/proteins were considered as true positive biomarkers; similarly, the same number of DEGs with the lowest fold changes were selected as true negative biomarkers (**Figure 2c-d**). The combined true positive and negative lists were input into ExIR as the desired lists of genes, and the outputs were compared with four commonly used biomarker prioritisation methods: mutual information (MI) ^27^, Student’s t-test, the point-biserial correlation coefficient, and the Spearman correlation coefficient. The evaluation and comparison of biomarker prioritisation methods were performed based on ROC analyses. MI was calculated between the expression profile of each gene and the binary (0,1) sample labels using the mpmi R package (https://cran.r-project.org/package=mpmi) ^77^. Similarly, point-biserial and Spearman correlation coefficients, as well as the Student’s t-test, were computed between gene expression profiles and binary sample labels using the stats R package. Additionally, immunohistochemical data deposited in the Human Protein Atlas was present for the exact lung adenocarcinoma (LUAD) subtype used for biomarker evaluation; thus, the expression of the top five ExIR-prioritised LUAD biomarkers in unaffected and LUAD tissue was examined using this resource ^28^ (http://www.proteinatlas.org). To further evaluate the potential of ExIR in identification and prioritisation of candidate biomarkers, the model was applied to the whole LUAD dataset without prior provision of a true positive or negative gene set and the immunohistochemical data of the Human Protein Atlas corresponding to the top five inferred biomarkers was examined in unaffected and LUAD samples.

### Evaluation of mediator prioritisation

As there exists no centralised resource containing validated sets of mediators of biological processes or diseases, the performance of ExIR in identifying and prioritising mediators was evaluated based on functional annotation of ExIR outputs. Initially, the performance of ExIR was assessed in comparison to the mediator genes inferred by the MALANI algorithm ^29^. In the context of cancer datasets, MALANI proposes two classes of genes; one being genes frequently differentially expressed or mutated, and another being those that are not differentially expressed but may mediate the coordination of oncogenic signals between DE/mutated genes ^29^. A set of breast cancer mediator genes has been proposed based on application of MALANI to the TCGA BRCA dataset; thus, to compare the performance of ExIR against MALANI for mediator detection, the entire TCGA BRCA dataset was input to ExIR without prior provision of any desired gene list. Next, an overrepresentation analysis (ORA) of all ExIR- and MALANI-derived mediators for biological processes and KEGG pathways was performed using the enrichR R package ^78^, and statistically non-significant terms were filtered out. The association of significant biological processes and KEGG pathways corresponding to ExIR- and MALANI-derived mediators in breast cancer were then separately interrogated using the Comparative Toxicogenomics Database (CTD, accessed October 19, 2021) ^79^, a manually-curated repository for literature-based and computationally inferred associations between genes, phenotypes, diseases, etc. Additionally, this benchmarking workflow was then applied beyond breast cancer to all other examined datasets.

### Animal husbandry

All protocols and procedures using zebrafish greater than 7 dpf were approved by the Monash University Animal Ethics Committee (ERM14481, ERM22161 and ERM17993). Zebrafish were maintained under standard housing and breeding conditions ^80^ in the AquaCore facility, Monash University. Embryos and larvae were maintained in E3 medium, while adult zebrafish were maintained in system water. Mutant strains used in this study were *sgsh^mnu301^* (referred to as *sgsh*) ^8^; transgenic lines used were *Tg*(*-8.7elavl3:Hsa.H2B-GCaMP6s)^jf5^;mitfa^w2/w2^*^81^.

### RNA sequencing

For each experimental replicate, total RNA was extracted from freshly-dissected n=3 young (3-month-old) or n=3 aged (18-month-old) zebrafish brain in TRIzol (Invitrogen, 15596). Three independent experimental replicates were used for bulk RNA sequencing, and all samples were assayed for RNA integrity on an Agilent 2100 Bioanalyzer using the Agilent RNA 6000 Nano Kit. 150 bp paired-end sequencing was performed by BGI (Hong Kong) using the DNBseq platform. SOAPnuke ^82^ was used for adaptor removal and low-quality read filtration, and genome mapping was performed with HISAT2 ^83^ for SNP analysis and novel transcript detection. Clean reads were mapped to the reference genome using Bowtie2 ^84^, and gene expression levels were calculated with RSEM ^85^. DESeq2 was used for differential gene expression analysis ^86^. Downstream visualisations for RNA-seq data were generated in R using EnhancedVolcano ^87^, GOplot ^88^ and ExIR (this paper), clusterProfiler ^89^, and in CytoScape ^90^ with the StringApp plugin^91^. DESeq2 differential expression analyses are provided in **Extended Data Tables 7-8**, and GO enrichment analyses of RNA datasets are listed in Extended Data Table 9.

### Proteomics

Label-based quantitation by tandem mass tagging (TMTpro 16plex, Thermo Scientific, A44520) of proteins was performed using nanoLC ESI MS/MS. n=5 samples were used per group. Total protein was purified from brains in 5% SDS with 100 mM HEPES at 95 °C for 10 minutes, then sonicated. Protein concentration was determined by BCA assay. Reduction and alkylation were performed using TCEP/CAA at 55 °C for 15 minutes, then protein was acidified using 1.2% phosphoric acid and captured with S-Trap spin columns (Protifi). Peptides were derived on-column using trypsin and Lys-C protease digestion. Following elution of peptides with 50 mM TEAB, columns were washed with 0.2% formic acid and then with 50% ACN with 0.2% formic acid to aid in recovery of hydrophobic peptide fragments. NanoLC ESI MS/MS was performed using a Dionex UltiMate 3000 RSLCnano (Thermo Scientific) and an Orbitrap Eclipse Tribrid mass spectrometer (Thermo Scientific). Analytical columns used were Acclaim PepMap RSLC (75 μm x 50 cm, nanoViper, C18, 2 μm, 100 Å, Thermo Scientific), and the trap column used was Acclaim PepMap 100 (100 μm x 2 cm, nanoViper, C18, 5 μm, 100 Å, Thermo Scientific). Initial data analysis was performed using Proteome Discoverer (Thermo Scientific) and SEQUEST with MS3 quantitation. The protein FDR cut-off was set at 1%, with fixed modifications of C-terminal carbamidomethylation and N-terminal TMTpro and variable modification of M oxidation and N-terminal acetylation. Subsequent data analysis was performed in R; data were filtered for high-confidence and contaminant proteins, and those proteins with a high proportion of missing values between samples were excluded. Protein intensity data was converted to a log2 scale, then samples were grouped by condition and missing values were imputed using the Missing Not At Random (MNAR) method which uses random draws from a left-shifted Gaussian distribution of 1.8 standard deviations apart with a width of 0.3. Data were normalised using the variance stabilising normalisation (vsn) method. Protein-wise linear models combined with empirical Bayes statistics were used for differential abundance analyses. limma ^68^ was used to generate a list of differentially abundant proteins for each pairwise comparison. A cut-off of the Benjamini-Hochberg adjusted *p*-value of 0.05 was applied, alongside a log2 fold-change cut-off of 1 to determine significantly differentially abundant proteins to account for the ratio compression of TMT-MS3 reporter usage. Downstream visualisations for proteomics data were generated in R using EnhancedVolcano ^87^, pathfindR ^92^ and ExIR (this paper), clusterProfiler ^89^, and CytoScape ^90^ with the StringApp plugin ^91^. Differential protein abundance analyses are provided in **Extended Data Table 11**, and GO enrichment analyses of proteomics datasets are listed in Extended Data Table 10.

### Real-time quantitative PCR (RT-qPCR)

Total RNA was extracted from samples using the TRIzol method and approximate concentration determined on a NanoDrop 1000 (Thermo Fisher Scientific). 1 μg total RNA was used for reverse transcription to cDNA using SuperScript IV reverse transcriptase (Invitrogen, 18090010) and a 1:1 mix of random hexamers and Oligo-dT(12-18). RT-qPCR was performed on a LightCycler 480 II (Roche) using LightCycler 480 SYBR Green I Master mix (Roche, 04707516001) using 1:20 cDNA dilution from stock. All primer pairs were tested for amplification efficiency between 90-110%. Relative gene expression was calculated in Excel (Microsoft) using the 2^-ΔΔCt^ method, with error propagation calculated as previously described ^93^. Primer sequences are listed in **Extended Data Table 5**.

### *Immunohistochemistry and* in situ *hybridisation*

Riboprobes for *in situ* hybridisation were generated by cloning transcript-specific PCR products into pGEM-T-Easy (Promega, A1360) using the TA-cloning method. Insert directionality was confirmed by Sanger sequencing, and plasmids were linearised to facilitate *in vitro* transcription of antisense riboprobes using either SP6 or T7 RNA polymerase and DIG-RNA labelling mix (Roche, 11277073910). All primers used to clone riboprobe sequences are listed in **Extended Data Table 5**.

Adult zebrafish were rapidly euthanised in an ice-water slurry, and exsanguinated on ice via a tail snip. Whole brains were dissected from the neurocranium in 1x phosphate-buffered saline (PBS) pH 7.4, and immediately transferred to 4% paraformaldehyde (PFA, Sigma, 158127) in PBS for overnight fixation at 4 °C with gentle rocking. After fixation, brains were cryoprotected in a sucrose-EDTA solution (20% sucrose, 20% 0.5 M EDTA pH 8 in 1x PBS) overnight at 4 °C, then cryo-embedded in a mixture of sucrose and fish gelatin as previously described ^94^. Brains were serially cryosectioned at 16 μm thickness using a Leica CS3050S cryostat.

For immunohistochemistry, samples were dried for >1 hour at room temperature (RT), then rehydrated with 1x PBS for 15 minutes. Sections were permeabilised 2x 15 minutes with 1x PBS + 0.3% Triton X-100 (PBS-Tx 0.3%) at RT, and incubated flat in a humified chamber with primary antibodies overnight at 4 °C. Sections were then washed 3x 20 minutes with PBS-Tx 0.3% at RT, then secondary antibodies were applied for one hour at RT at indicated concentrations with 5 mg/mL DAPI (Sigma, D9542) applied at 1:5000 concentration as nuclear counterstain. A single 10-minute PBS-Tx 0.3% wash was then performed followed by 2x 20-minute washes with 1x PBS, mounted with 50% glycerol and coverslipped prior to imaging. Sections were imaged on a Leica TCS SP8 confocal microscope equipped with a HyD detector. All antibodies used and the concentrations used are listed in **Extended Data Table 6**.

For *in situ* hybridisation, sections were pre-fixed with 4% PFA in PBS pH 7.4 for one hour, then washed twice with PBS-Tx 0.3% for 20 minutes. 10 mg/mL Proteinase K (Roche, 3115879001) was diluted 1:500 in PBS-Tx 0.3%, and sections were digested at RT for five minutes. Sections were then quickly washed with PBS-Tx, and Proteinase K digestion was stopped by incubating sections with 4% PFA at RT for 10 minutes, followed by 2x 10-minute washes with PBS-Tx 0.3%. A hybridisation chamber was assembled using a slide box, containing filter paper saturated with a hybridisation chamber solution (5 mL 10x Salt solution [1.95 M NaCl, 89 mM Tris-HCl, 11 mM Tris base, 50 mM NaH2PO4·2H2O, 50 mM Na2HPO4 and 63.68 mM EDTA], 25 mL formamide and 20 mL ddH2O, and the hybridisation chamber was preheated to 60 °C in an incubator. Antisense riboprobes were diluted 1:200 in hybridisation buffer (1 mg/mL Torula RNA, 50% formamide, 1x Salt solution, 10% dextran sulfate, 1x Denhardt’s buffer, in ddH2O), vortexed and denatured at 70 °C for 10 minutes prior to addition to sections. Parafilm was used to mitigate probe evaporation during hybridisation. Probes were hybridised overnight at 60 °C. Unbound probe was then removed by washing with 1x SSC buffer and 50% formamide in ddH2O 1x 15 minutes, then 2x 30 minutes at 62 °C, followed by 2x 30-minute washes at RT in MABT. Sections were blocked in 2% DIG blocking reagent (Roche, 11096176001) in MABT for two hours at RT, then incubated for four hours at RT in AP-conjugated anti-DIG Fab fragments (Roche, 11093274910) diluted 1:2000 in 2% DIG blocking reagent. Sections were then washed 4x 20 minutes in MABT at RT, and equilibrated in staining buffer (0.1 M NaCl, 0.05 M MgCl2, 0.1 M Tris pH 9.5, Tween-20 at 1:1000, all in ddH2O) for 5 minutes. Chromogenic detection was then performed using NBT-BCIP stock solution (Roche, 11681451001) diluted 1:50 in staining buffer until sufficient signal was observed. Development was terminated by incubating sections in 4% PFA for 30 minutes, then washed 3x 10 minutes in PBS. Sections were mounted with 50% glycerol prior to imaging on an Olympus IX-81 inverted microscope.

### Transmission electron microscopy

Adult zebrafish were euthanised and brains extracted from the neurocranium as above, then sub-dissected into separate forebrain, midbrain and hindbrain components. Brain regions were immediately transferred into Karnovsky fixative (2.5% glutaraldehyde and 2% PFA, with 0.25% CaCl_2_ and 0.25% MgCl_2_ in 0.1 M sodium cacodylate, pH 7.4) and fixed at RT for 2 hours, then post-fixed in 1% osmium tetroxide/1.5% potassium ferricyanide followed by 5× 10-minute washes in distilled water. Tissue was then incubated in 70% ethanol overnight, then dehydrated in a stepwise, increasing ethanol gradient (2× 10-minute washes in 80%, 90% and 95% ethanol, followed by 4× 10-minute washes in 100% ethanol). Samples were embedded in Epon EMbed 812 resin (Electron Microscopy Sciences, 14120). Ultrathin (70 nm) sections were obtained using a Leica UltraCut UC7 and stained with uranyl acetate and lead citrate. Sections were imaged at 80 kV on a Jeol 1400+ transmission electron microscope.

### In vivo *neuronal calcium imaging in zebrafish*

*In vivo* calcium imaging was performed as previously described ^95^. Briefly, transgenic *Tg*(*elavl3 :Hsa.H2B-GCaMP6s*)^*jf5*^;mitfa^*w2/w2*^;sgsh^*mnu301/mnu301*^ or *Tg*(*elavl3:Hsa.H2B-GCaMP6s*)^*jf5*^;mitfa^*w2/w2*^;sgsh^*WT/WT*^ larvae at 7 dpf were anaesthetised using 0.03% tricaine methanesulfonate and mounted in a droplet of 0.5% low-melting temperature agarose (Sigma, A9414) in Ringer’s solution pH 7 (116 mM NaCl, 2.9 mM KCl, 1.8 mM CaCl_2_, 5 mM HEPES in ddH_2_O) on a glass slide. When the agarose droplet solidified, a plastic ring was sealed to the slide around the droplet with a thin application of petroleum jelly to form a water-tight immersion chamber. 1 mL of Ringer’s solution was applied to the immersion chamber and larvae were monitored for recovery of spontaneous motor function. Larvae were then imaged for 5 minutes on a Zeiss Axio Imager Z1 with a Zeiss EC Plan-NEOFLUAR 5x 0.16 NA objective, with images captured every 0.5 seconds to establish baseline GCaMP6s activity. Following this, pentylenetetrazole (PTZ, Sigma, P6500) was added to the immersion chamber to a final concentration of 20 mM ^96^, and larvae were incubated in darkness for three minutes then imaged for a further two five minute intervals. Time-lapse acquisitions were exported to ImageJ2 ^97^, and the ‘Set Measurements’ option was employed to derive the mean grey value from image data. Using the polygon selection tool, an ROI was drawn around the whole brain, and stored in the ROI manager. The ‘Multi Measure’ functionality was then used to measure the mean grey value of the ROI in all slices in the time-lapse. These data were exported to GraphPad Prism 9 (GraphPad Software) for downstream analysis.

## Data and code availability

All analyses used for the evaluation and benchmarking of ExIR were performed on publicly available data. The TCGA datasets are available at https://gdac.broadinstitute.org. Raw and normalised GBM scRNA-seq data were downloaded from http://www.gbmseq.org, which is also available from the GEO database under the accession number GSE84465. Raw and normalised oogenesis scRNA-seq data ^25^ were retrieved from GEO accession GSE130212. Schizophrenia microarray data ^21^ were retrieved from GEO accession GSE93577. The software and source code for the ExIR model is available as a function in the influential R package, accessible at https://github.com/asalavaty/influential or from the CRAN repository (https://cran.r-project.org/package=influential). Instructions on how to prepare input data for ExIR and how to execute functions, as well as some examples based on simulated data are available at https://cran.r-project.org/web/packages/influential/vignettes/Vignettes.html. Any requests for additional information will be fulfilled by the lead contacts. All transcriptomic and proteomic datasets generated in this study will be uploaded to the NCBI Gene Expression Omnibus and the Proteomics Identification Database (PRIDE), respectively, prior to publication.

## Acknowledgements

The authors would like to thank Lan Nguyen for their constructive feedback on the development of ExIR and proteomic data analysis, and Monash AquaCore, and Monash Micro Imaging facilities for their excellent support. The authors acknowledge the use of instruments and assistance at the Monash Ramaciotti Centre for Cryo-Electron Microscopy, a Node of Microscopy Australia. This study also used infrastructure located at the Monash Proteomics and Metabolomics Facility, enabled by Bioplatforms Australia (BPA) and the National Collaborative Research Infrastructure Strategy (NCRIS). The results shown in this study are in part based on data generated by the TCGA Research Network (http://cancergenome.nih.gov/). This work was supported by Monash University, a National Health and Medical Research Council (NHMRC) Fellowship (1136567) to PDC, an NHMRC grant (1180905) to MR, an incubator grant from the Sanfilippo Children’s Foundation (Australia) and the Cure Sanfilippo Foundation (US) to JK, a Translational grant from the Sanfilippo Children’s Foundation (Australia), the Cure Sanfilippo Foundation (US) and Fundacja Sanfilippo (Poland) to JK. AD, AS, FK and S-AS are supported by Australian Government Research Training Program (RTP) scholarships. The Australian Regenerative Medicine Institute is supported by grants from the State Government of Victoria and the Australian Government.

## Author contributions

AD, AS, MR, PDC and JK conceptualised the study. AS developed ExIR and performed *in silico* benchmarking and data analysis. AD performed all *in vivo* experiments and analysis with contributions from FK and S-AS. AD, JRS, IH, ADS and RBS performed proteomics study design, sample preparation, mass spectrometric acquisition and preliminary data analysis. AD and AS wrote the manuscript with contributions from MR, PDC and JK.

## Figure legends

**Extended Data Fig. 1 | Non-telencephalic *baiap2b* expression and choroid plexus *pmela* expression in adult wild type and *sgsh* zebrafish brain. a-b’,***baiap2b* expression in midbrain was detected at comparable levels between wild type and *sgsh* mutants in the dorsal tegmental nucleus. **c-d’,** *baiap2b* expression in hindbrain was detected at comparable levels between wild type and *sgsh* mutants in neurons in cells in the Purkinje layer, presumably Purkinje or eurydendroid cells. **e,** *pmela* expression was not detected in wild type adult brain. **f,** *pmela* expression was detected in, and highly restricted to, the choroid plexus dorsal to the thalamus/preoptic region. Scale bars 100 μm in a, b, c, d, e, f; 40 μm in a’, b’, c’, d’.

**Extended Data Fig. 2 | ExIR outperforms other common feature prioritisation methods. a-b,** Criteria used to define ground truth true positive and negative driver gene sets for benchmarking in homeostatic (**a**) and disease datasets (**b**). Note the requirement for a training set required for execution of Endeavour and ToppGene, which is not required for driver gene determination using ExIR. **c-d**, Criteria used to define true positive and negative biomarkers from cancer datasets (**c**) and those of other diseases (**d**). **e**, Raincloud plot summaries of ROC plot AUCs for comparison of performance of driver gene prioritisation methods. **f**, Raincloud plot summaries of ROC plot AUCs for comparison of performance of biomarker gene prioritisation methods. **g,** ROC plots (true positive vs. false positive rates) of driver prioritisation performance in TCGA BRCA, LIHC, LUAD, THCA, and GBM (GSE130212), Schizophrenia (GSE93577), BRCA (PXD002057) and LUAD (IPX0001804000) datasets. **h,** ROC plots (true positive vs. false positive rates) of biomarker prioritisation performance in TCGA BRCA and LUAD, and Schizophrenia (GSE93577) datasets. **i,** Quantification of TCGA BRCA GO-BPs and KEGG pathways corresponding to ExIR- and MALANI-derived mediators. **j,** Quantification of ExIR- and MALANI-derived mediators enriched in TCGA BRCA GO-BPs and KEGG pathways. **k,** Number of associated GO-BP or KEGG pathway annotations associated with ExIR-derived mediators across different disease datasets. **l,** Percentage of ExIR-derived mediators enriched in disease-associated GO-BPs and KEGG pathways. TCGA, The Cancer Genome Atlas; GBM, glioblastoma multiforme; BRCA, breast cancer; LIHC, liver hepatocellular carcinoma; LUAD, lung adenocarcinoma; THCA, thyroid cancer.

**Extended Data Fig. 3 | Comparison of top five known and ExIR-predicted LUAD biomarkers. a-e,** Immunohistochemical (IHC) data from the Human Protein Atlas ^28^ database in LUAD and normal lung tissue for top five known LUAD biomarkers. **a,** SFTPC - ExIR rank #1, <u>LUAD</u> (negative intensity; patient id: 1847) and <u>normal pneumocytes</u> (quantity:75%-25%; strong intensity; patient id: 2268). **b,** SPP1 – ExIR rank #117, <u>LUAD</u> (quantity:>75%; moderate intensity; patient id: 537) and <u>normal pneumocytes</u> (not detected; patient id: 2268). **c,** CBLC – ExIR rank #140, <u>LUAD</u> (quantity:>75%; moderate intensity; patient id: 1847) and <u>normal pneumocytes</u> (not detected; patient id: 2417). **d,** MDK – ExIR rank #247, <u>LUAD</u> (quantity:75%-25%; strong intensity; patient id: 1847) and <u>normal pneumocytes</u> (not detected; patient id: 2222). **e,** MRC1 – ExIR rank #471, <u>LUAD</u> (negative intensity; patient id: 1932) and <u>normal macrophages</u> (quantity:75%-25%; strong intensity; patient id: 2208). **f-i,** IHC data of top five ExIR-predicted LUAD biomarkers (excluding SFTPC rank #1 already in **a**). **f,** AGER – ExIR rank #2, <u>LUAD</u> (not detected; patient id: 3144) and <u>normal pneumocytes</u> (quantity:>75%; strong intensity; patient id: 4840). **g,** EMP2 – ExIR rank #3, <u>LUAD</u> (not detected; patient id: 1847) and <u>normal pneumocytes</u> (quantity:>75%; strong intensity; patient id: 2101). **h,** CAV1 – ExIR rank #4, <u>LUAD</u> (not detected; patient id: 1249) and <u>normal pneumocytes</u> (quantity:>75%; strong intensity; patient id: 2208). **i,** RTKN2 – ExIR rank 5, <u>LUAD</u> (not detected; patient id: 3003) and <u>normal pneumocytes</u> (quantity: <25%; moderate intensity; patient id: 2268). As requested by The Human Protein Atlas database, the link to the immunostaining images of all of the selected proteins in normal pneumocytes and <u>LUAD</u> samples are included as hyperlinks within the figure legend. Ab, antibody; LUAD, lung adenocarcinoma.

**Extended Data Fig. 4 | Top-ranked driver and biomarker genes and proteins in *sgsh* mutant RNA-sequencing and quantitative proteomics datasets. a,** ExIR prioritisation of upregulated (accelerating) driver and biomarker genes amongst young brain differentially abundant genes (DEGs). **b,** ExIR prioritisation of downregulated (decelerating) driver and biomarker genes amongst young brain DEGs. **c,** ExIR prioritisation of upregulated (accelerating) driver and biomarker proteins amongst young brain differentially abundant proteins (DAPs). **d,** ExIR prioritisation of downregulated (decelerating) driver and biomarker proteins amongst young brain differentially abundant proteins (DAPs). **e,** ExIR prioritisation of upregulated (accelerating) driver and biomarker genes amongst aged brain DEGs. **f,** ExIR prioritisation of downregulated (decelerating) driver and biomarker genes amongst aged brain DEGs. **g,** ExIR prioritisation of upregulated (accelerating) driver and biomarker proteins amongst aged brain DAPs. **h,** ExIR prioritisation of downregulated (decelerating) driver and biomarker proteins amongst aged brain DAPs.

**Extended Data Fig. 5 | Functional enrichment analyses of cross-timepoint conserved transcriptomic and proteomic mediators. a,** STRING PPI network (interaction confidence cut-off 0.9, singletons not displayed) of ExIR-derived mediators present in both young and aged RNA-seq datasets. **b,** Tree plot of Gene Ontology (GO) functional enrichment analysis (biological process, molecular function and cell component) of conserved transcriptomic mediators. Node size is number of genes per GO term. Node colour is adjusted P-value *(Padj).* **c,** STRING PPI network (interaction confidence cut-off 0.9, singletons not displayed) of ExIR-derived mediators present in both young and aged proteomics datasets. **d,** Top 10 protein mediators in young *sgsh* brain. **e,** Top 10 protein mediators in aged *sgsh* brain. **f,** Tree plot of Gene Ontology (GO) functional enrichment analysis (biological process, molecular function and cell component) of conserved proteomic mediators. Node size is number of genes per GO term. Node colour is adjusted P-value (P*adj*).

**Extended Data Fig. 6 | Basal IEG expression is not different between wild type and *sgsh* mutants at 7 dpf. a,** RT-qPCR analysis of *egr1, egr4,* and *npas4a* expression (normalised to *eef1a1l1* expression) in dissected brains of wild type or *sgsh* 7 dpf larvae. Each group is RNA from n=20 pooled brains. *n.s.,* not significant. **b-c,** *egr1* expression detected by *in situ* hybridisation in 7 dpf larval brain in wild type and *sgsh* homozygotes. **d-e,** *egr4* expression detected by *in situ* hybridisation in 7 dpf larval brain in wild type and *sgsh* homozygotes. **f,** Experimental schematic for PTZ exposure and *npas4a* detection in 7 dpf larvae of either genotype. **g,** *npas4a* expression in wild type 7 dpf telencephalon (n=20) following 30 minutes of 20 mM PTZ exposure. Scale bar 50 μm. **h,** *npas4a* expression in *sgsh* 7 dpf telencephalon (n=20) following 30 minutes of 20 mM PTZ exposure. n=2 samples showed no expression, presumably due to failed hybridisation. Scale bar 50 μm. **i-k,** GCaMP6s fluorescence (whole brain) at baseline and after PTZ administration in n=3 *sgsh^WT^* 7 dpf larvae. **l-n,** GCaMP6s fluorescence (whole brain) at baseline and after PTZ administration in n=3 *sgsh* 7 dpf larvae.

**Extended Data Table 1 | Biological processes significantly associated with mediators of investigated diseases from ExIR and MALANI models.**

**Extended Data Table 2 | Disease-associated biological processes corresponding to ExIR-or MALANI-derived mediators.**

**Extended Data Table 3 | Curated ground-truth driver gene sets used for benchmarking.**

**Extended Data Table 4 | Curated ground-truth biomarker gene sets used for benchmarking.**

**Extended Data Table 5 | Primer sequences used for RT-qPCR and riboprobe cloning.**

**Extended Data Table 6 | Antibodies and dyes used for immunohistochemistry.**

**Extended Data Table 7 | DEseq2 differential expression analysis of young wild type and *sgsh* brain.**

**Extended Data Table 8 | DEseq2 differential expression analysis of aged wild type and *sgsh* brain.**

**Extended Data Table 9 | GO analyses of transcriptomic mediators conserved across ExIR analyses of young and aged timepoints.**

**Extended Data Table 10 | GO analyses of proteomic mediators conserved across ExIR analyses of young and aged timepoints.**

**Extended Data Table 11 | Differential abundance analyses of young and aged wild type and *sgsh* brain proteomics.**

## Notes

### Competing Interest Statement

The authors have declared no competing interest.

